# The Cellular Underpinnings of the Human Cortical Connectome

**DOI:** 10.1101/2023.07.05.547828

**Authors:** Xi-Han Zhang, Kevin M. Anderson, Hao-Ming Dong, Sidhant Chopra, Elvisha Dhamala, Prashant S. Emani, Daniel Margulies, Avram J. Holmes

## Abstract

The functional properties of the human brain arise, in part, from the vast assortment of cell types that pattern the cortex. The cortical sheet can be broadly divided into distinct networks, which are further embedded into processing streams, or gradients, that extend from unimodal systems through higher-order association territories. Here, using transcriptional data from the Allen Human Brain Atlas, we demonstrate that imputed cell type distributions are spatially coupled to the functional organization of cortex, as estimated through fMRI. Cortical cellular profiles follow the macro-scale organization of the functional gradients as well as the associated large-scale networks. Distinct cellular fingerprints were evident across networks, and a classifier trained on post-mortem cell-type distributions was able to predict the functional network allegiance of cortical tissue samples. These data indicate that the *in vivo* organization of the cortical sheet is reflected in the spatial variability of its cellular composition.

## Introduction

A core goal of research in the brain sciences is to understand the multiscale relationships that link molecular and cellular processes with the *in vivo* functional organization of the human cerebral cortex. Historically, the localization of the borders and associated areal parcels along the cortical sheet were determined by invasive techniques including histology, anatomical tract tracing, electrophysiology, and lesion methods. Through these approaches neuroscientists and histologists produced landmark maps that divide cortical territories on the basis of regional patterns of cytoarchitecture^1–8^, revealing the presence of both serial and parallel information processing hierarchies^9,10^. Recently, the development of dense spatial transcriptional atlases has enabled the study of cellular correlates of brain functions in humans, for instance as estimated through functional magnetic resonance imaging (fMRI)^11^. Initial work in this area has established molecular correlates of large-scale functional network organization^11–13^, including genes encoding ion channels^14^ and those enriched in supragranular layers of cortex^15^, as well as associations between the spatial distribution of interneuron-linked genes and regional differences in fMRI signal variability^16,17^. However, the extent to which associated *ex vivo* cellular architectures may mirror the hierarchical functional properties of the human cerebral cortex as measured by resting state fMRI (rs-fMRI) has yet to be systematically investigated.

From sensation through cognition and action, the human cortex is organized into a multiscale system composed of areal units that are situated along segregated processing streams^10^. These areal units, or parcels, are embedded within corresponding large-scale functional networks that are evident through both anatomical projections, task-evoked activity, and patterns of coherent neural activity at rest^18–20^. Supporting this network architecture, converging evidence indicates the presence of a broad division separating unimodal somatosensory/motor (somato/motor) and visual territories from the heteromodal association areas that integrate long-distance projections across distributed brain systems^9,21^. This hierarchical property of brain organization is reflected in the presence of functional gradients that span the cortical sheet, situating functionally distinct networks and corresponding areal parcels along a continuous spectrum^22^. These gradients reflect low-dimensional representations of functional connectivity, with the first, or primary, gradient anchored at one end by unimodal regions supporting primary sensory or motor functions and at the other end by the association cortex. The second gradient of connectivity reflects a sensory organization (unimodal gradient) anchored at each end by either visual or somato/motor cortex^22,23^. However, while recent evidence suggests a genetic basis for the macro-scale organization of the cortical sheet^24–27^, the extent to which cellular processes may underpin the functional organization of the brain remain to be established.

The translational challenge of linking molecular and cellular processes with properties of functional organization is addressable, in part, by integrating transcriptional data from *ex vivo* tissue samples with estimates of *in vivo* brain function^11^. Classic neuroanatomical discoveries revealed the evolutionary processes and developmental mechanisms that constrain the layout of cortical areas, their corresponding microstructure, and anatomical connectivity^28–30^. Recent work supports the presence of broad axes of cortical organization^30,31^, for instance as reflected in the spatial distribution of receptor densities^32^, intracortical myelination^33^, and supragranular-infragranular pyramidal neuron soma size ratios^34^. Preliminary studies associating gene expression to functional networks have revealed shared enrichment of genes among anatomical regions that are functionally coupled^12,14,15^, perhaps indicating the network-preferential presence of particular cell types. In line with this conjecture, subsequent work has revealed that spatial profiles of signal variability observable in BOLD fMRI follow the relative distribution of certain classes of inhibitory interneurons, for instance parvalbumin (PVALB) and somatostatin (SST)^16^, and broadly separate unimodal and association cortices. Intriguingly, select aerial boundaries derived from rs-fMRI have also been shown to correspond to histologically and structurally defined architectonic areas^19,23,35,36^. However, while these data suggest a link between the micro-(molecular and cellular) and macro-scale (gradients and networks) properties of brain organization, our understanding of how the complex functional architecture of the cerebral cortex develops and is maintained over the lifespan remains fragmentary. One possibility is that the relative preponderance of certain cell classes may spatially couple to gradually shifting gradient patterns across the cortical sheet. An alternative, but not mutually exclusive, hypothesis is that the spatial distribution of cortical cell types tracks the topographic organization of functionally connected but spatially distinct large-scale cortical networks.

Here, we examine the association between cortical gradients, functional networks, and the spatial distribution of cortical cell types, inferred from patterns of gene transcription in bulk tissue data. In doing so, we demonstrate that imputed cell type distributions spatially track the macro-scale gradient organization of cortex, both at the level of individual cell types and multivariate cellular profiles. Suggesting the presence of a complementary network structure of cellular organization, distinct cellular enrichment patterns were also evident across large-scale cortical networks. These “cellular fingerprints” can be used to predict the network allegiance of cortical parcels from their corresponding cell type abundance measured in independent post-mortem brain tissue. These data help address a key challenge in neuroscience to understand how cell-type distributions may underlie the *in vivo* functional properties of the human brain, establishing spatial correspondence between regional cellular profiles and the hierarchical organization of the cortical sheet.

## Results

### The large scale functional organization of human cortex

The multiscale and hierarchical organization of cortical functions were characterized by macroscale gradients and large-scale networks (Fig. 1). Here, we applied the diffusion map embedding^22,37,38^ to decompose vertex-level rs-fMRI functional connectivity (FC) matrices into continuous gradients that capture the maximum variance across the 820 adults in the Human Connectome Project (HCP)^39^. To associate the functional measures with cellular profiles, the gradient values were parceled into the Schaefer-400 atlas^35^. Parcel-level network assignment was obtained as detailed in Yeo et al. 2011^23^ (Fig. 1A). Next, we use these complementary gradient-and network-based approaches as a foundation to establish the cellular associates of the *in vivo* functional organization of the cerebral cortex.

**Fig. 1.**
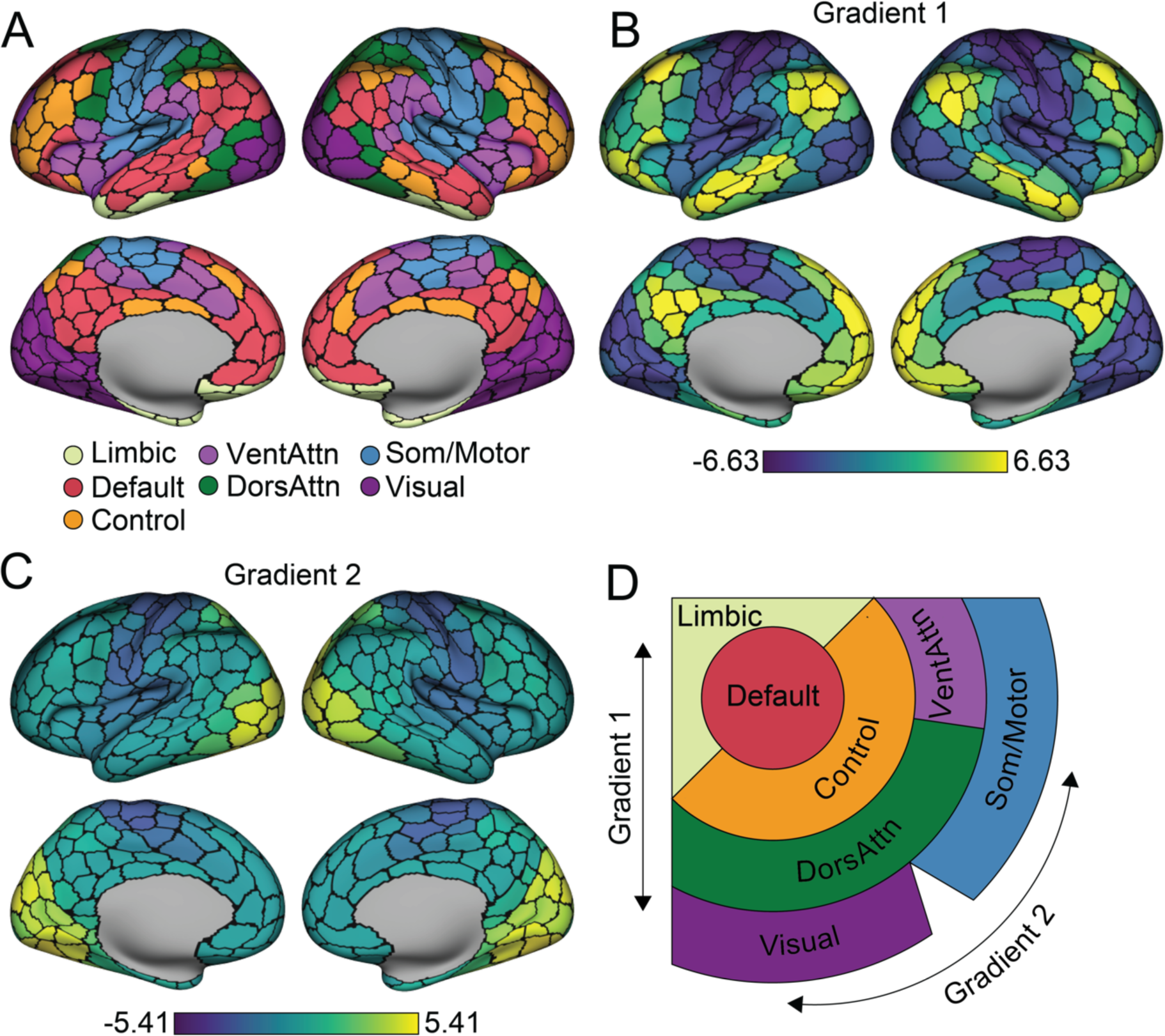
Large-scale functional networks are embedded along two principal gradients. **A.** Functionally coupled cortical parcels are grouped into large-scale networks, based on the Yeo et al.^23^ 7-network solution averaged across the 400-parcel functional atlas of Schaefer et al.^35^ **B.** The first principal gradient of intrinsic functional connectivity. Parcels are colored by their relative topological position spanning between the association cortex (bright yellow) and the unimodal cortex (dark blue). Scale bar reflects z-transformed principal gradient values derived from connectivity matrices using diffusion map embedding^37^. **C.** The second gradient of intrinsic functional connectivity is anchored within unimodal areas including primary visual cortex (bright yellow) at one end and somatomotor/auditory cortex at the other (dark blue). **D**. Figure displays the spatial organization of the seven networks along the two primary gradients. Adapted from Margulies et al.^22^. VentAttn, salience/ventral attention network; DorsAttn, dorsal attention network; Som/Motor, somato/motor network.

The current analyses focused on the first two primary gradients, reflecting the canonical information processing hierarchies in the human cortex^10^. Unlike linear methods that reduce geometric dimensionality, diffusion map embedding allows topologically similar local and long distance connections to be placed into common spaces with interpretable architectures^22,38,40^. The resulting gradients are unitless and reflect the position of vertices along an associated embedding axis that captures the primary differences in FC patterns. Consistent with prior work^22,38^, the architecture of the first gradient (Gradient 1) spans from unimodal areas (including primary visual, auditory, somatosensory, and motor cortex) through transmodal, or association (default network), territories (Fig. 1B). The peak values in the second gradient (Gradient 2) were evident along the central and calcarine sulcus, differentiating the somato/motor cortex from the primary visual system (Fig. 1C). The first two gradients account for a substantial proportion of the variance in functional connectivity (Gradient 1: 26%; Gradient 2: 12%). As initially reported by Margulies et al.^22^, large-scale functional networks^23,35^ are distributed across the cortical sheet and spatially ordered along these first two primary gradients (Fig. 1D), a property of cortical organization reflected in the repeating transitions between networks across cortical lobes. These data highlight complementary analytic frameworks that situate large-scale cortical networks and functions in separate domains along overlapping organizing axes.

### Univariate cellular associates of cortical gradients

Areal parcellations derived from rs-fMRI have been shown to follow boundaries of select histologically- and structurally-defined architectonic areas, for instance within somato/motor territories and putative language areas^23,35^. However, the extent to which regional cellular profiles spatially covary with the broad hierarchical organization of cortex has yet to be fully established *in vivo*. To examine the relationship between the topographic distribution of individual *ex vivo* cell types and the two primary functional gradients, the cortical topography of cell type abundances was inferred from Allen Human Brain Atlas (AHBA) post-mortem bulk gene expression samples using a previously validated method (see Methods)^11^. The transcription signatures identifying each class of cell in the AHBA bulk samples were derived from cortical single-nucleus droplet-based sequencing (snDrop-seq) data of visual and frontal cortex as reported by Lake et. al.^41^. The snDrop-seq data from the two regions allowed for parallel validation of the imputation process. The molecular signature profiles of 17 cell classes (see Methods) were constructed from snDrop-seq samples, including 5 interneuron subtypes (i.e., PVALB, SST, In1, In3, and In4), 6 excitatory neuron subtypes (i.e., Ex1, Ex3, Ex4, Ex5, Ex6, and Ex8), and 6 non-neuronal subtypes (astrocytes, oligodendrocytes, pericytes, endothelial, microglia, and oligodendrocyte precursor cells). The abundance of each cell type was estimated in available bulk samples which were further aggregated into the 400 cortical parcels (see Methods). The resulting distribution of parcel-level cell type abundances were examined relative to the *in vivo* functional gradient organization of the cortex. Statistical significance was established using permuted spin tests accounting for the spatial autocorrelation^42,43^, FDR corrected for 34 multiple comparisons (17 cell types x 2 gradients).

The association between the two gradients and the 17 cell types were separately imputed using visual (Lake VIS) and frontal cortex (Lake DFC) samples are displayed in Fig. 2A. To minimize the effects of spatial heterogeneity of single-cell transcriptional signatures, we only discuss subtypes that survived multiple comparison correction across both Lake DFC and Lake VIS imputed samples and were therefore robust in their associations with gradient architecture. Fig. 2B-D shows the results based on Lake DFC, and the full results are displayed in Supplementary Fig 1-2. The first gradient (Gradient 1; Fig. 2B), spanning the unimodal-transmodal axis, was positively associated with the imputed spatial distributions of interneuron In1 (Lake DFC: Spearman correlation, Rho=0.309, p_FDR_= 0.011; Lake VIS: Rho=0.290, p_FDR_=0.041) and excitatory neuron Ex1 cells (Lake DFC: Rho=0.307, p_FDR_<0.001; Lake VIS: Rho=0.314, p_FDR_=0.009). Gene markers of both of these cell types are primarily clustered in the molecular layer of cortex (layer I)^41^.

**Fig. 2.**
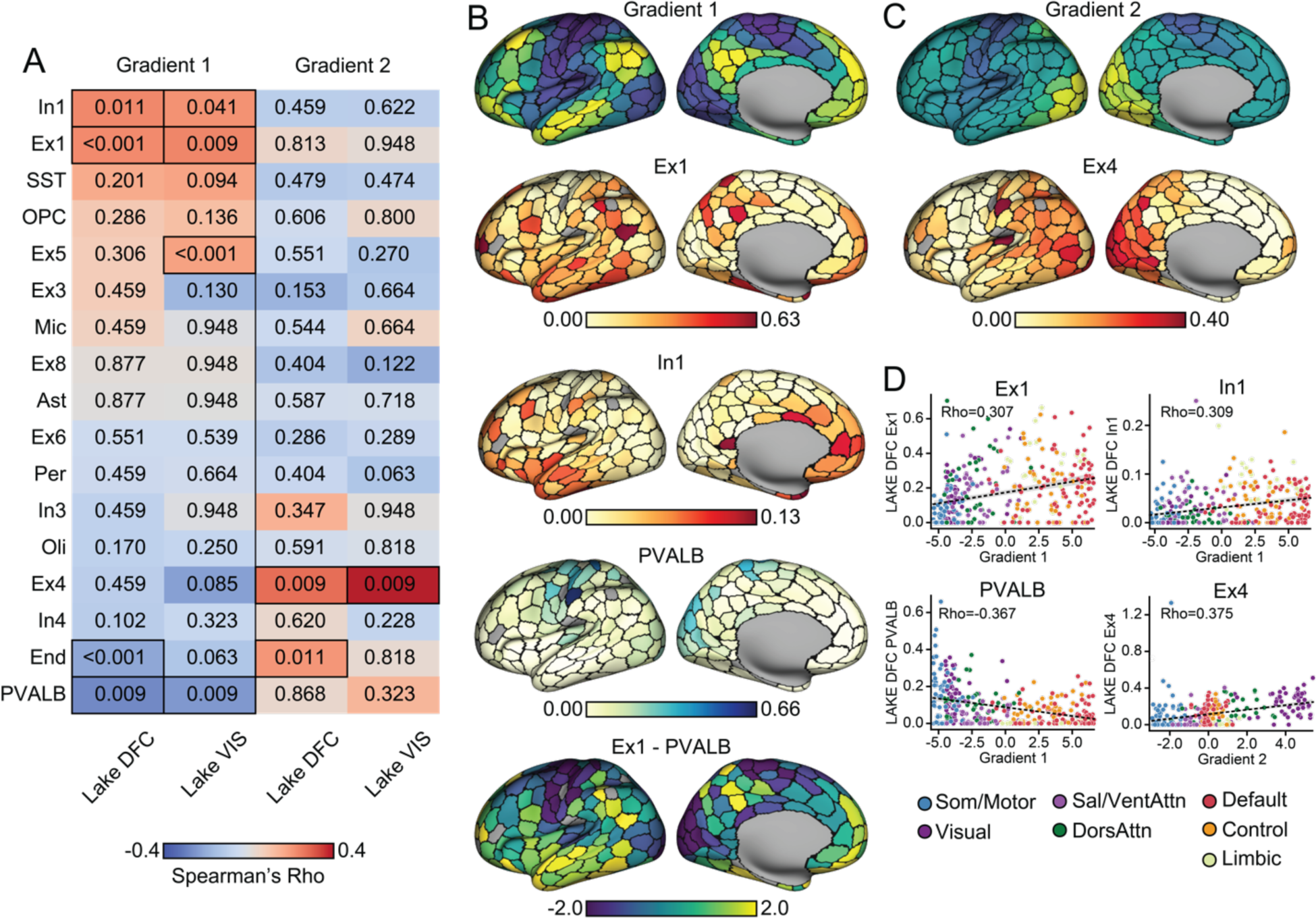
Univariate cell type distributions align with functional gradient topographies. **A**. Cell types are imputed from gene expression in AHBA bulk tissue samples. Single-cell signatures are constructed from independent tissue samples in the frontal (Lake DFC) and visual (Lake VIS) lobes^41^, allowing for a technical replication of the cell-type imputation scheme. Resulting abundance of cell types were rank-ordered by spatial correlation to each principal functional gradient. Warm colors indicate positive correlation, and numbers in each cell reflect associated FDR corrected p_spin_. Significant correlations were boxed in **A**. Surviving from significance tests in both Lake DFC and Lake VIS, the spatial pattern of Gradient 1 was correlated with two interneuron subtypes: In1 and PVALB, as well as one excitatory neuron subtype: Ex1. Gradient 2 was significantly correlated to Ex4 excitatory neurons. **B.** Imputed cell type abundance distributed across cortex suggests Ex1 and In1 are preferably distributed around the transmodal end of Gradient 1 (bright yellow), whereas PVALB is preferably distributed around the unimodal end (dark blue). A difference score between Ex1 and PVALB distributions generates a pattern spatially consistent with the first functional gradient. **C.** Ex4 follows a spatial pattern aligning to the second gradient, peaking in visual pole (bright yellow) then gradually decreasing as it approaches the somato/motor and auditory cortices (dark blue). Parcels that are excluded from analyses, not covered by AHBA bulk samples, are colored in gray. **D.** Scatter plots with each cortical parcel colored by the corresponding functional networks show that cell type abundance gradually increase/decrease across the networks distributed along the gradients, with enrichment/absence evident within certain networks. Correlations were estimated by Spearman’s Rho (as reflected in the scale bar in **A**), but for visual reference, dotted lines reflecting linear correlation between cellular abundance and gradient values are also displayed.

As shown in Fig. 2B, Ex1 is densely distributed within: 1) temporoparietal junction, posteromedial cortex, superior frontal gyrus, superior temporal sulcus in the default network; 2) inferior temporal gyrus, inferior frontal gyrus in the dorsal attention network; and 3) select aspects of the frontoparietal network, including the frontal pole and inferior parietal lobe. Ex1 cells (formally CBLN2^+^RASGRF2^+^ Ex1)^41^ are a subset of excitatory neurons marked by high expression of cerebellin 2 precursor (CBLN2) and ras protein specific guanine nucleotide releasing factor 2 (RASGRF2). CBLN2 is an extracellular synapses organizer^44–46^ particularly relevant to the expansion of dendritic spines within the human prefrontal cortex, compared to non-human primate^47^. RASGRF2 contributes to synaptic plasticity by mediating the induction of long-term potentiation^48,49^. In1 (including CCK^+^CNR1^+^RELN^+^ In1a, CCK^+^CNR1^+^THSD7B^+^ In1b, and VIP^+^CALB2^+^TAC3^+^ In1c)^41^ are a group of interneurons that exhibit a high expression of cholecystokinin (CCK), cannabinoid receptor 1 (CNR1). Of note, this class of cells includes subtypes like VIP and RELN inhibitory interneurons. As such, In1 cells can be marked by high expression of Reelin (RELN), thrombospondin type 1 domain containing 7B (THSD7B) or vasoactive intestinal peptide (VIP). In the current data, In1 peaks around the frontal and temporal pole, overlapping with default, ventral attention and dorsal attention network parcels (Fig 2B). In contrast, a negative association was observed between PVALB interneurons (Lake DFC: Rho=-0.367, p_FDR_=0.009; Lake VIS: Rho=-0.325, p_FDR_=0.009) and the first gradient. Consistent with prior work^16,17^, PVALB inhibitory interneurons, whose markers preferentially cluster in internal granule cell layer (layer IV)^41^, are preferentially distributed in somato/motor and visual areas such as postcentral gyrus, paracentral lobule and medial occipital lobe (Fig. 2B).

When considering Gradient 2, the analyses revealed a positive spatial association with excitatory Ex4 cells (formally RORB^+^IL1RAPL2^+^TSHZ2^+^FOXP2^+^; Lake DFC: Rho=0.375, p_FDR_<0.001; Lake VIS: Rho=0.535; p_FDR_<0.001; Fig 2C). Here, Ex4 presented a spectrum that peaked at visual and dorsal attention network parcels then gradually decreased along the dorsal attention network through default network areas, with increased presence within a subset of somato/motor parcels. The estimated preponderances of imputed cell types are displayed across cortical parcels (Gradient 1, Fig. 2B; Gradient 2, Fig. 2C), demonstrating that cell type abundance gradually increase/decrease across the functional networks distributed along the gradients, with their relative enrichment or absence evident within certain networks (Fig. 2D). The value of cell type abundance is always positive and gradient values represent the parcels’ topological position relative to each other. Accordingly, both positive and negative correlations indicate that the presence of a given cell type follows the corresponding gradient values. Of note, while these analyses reveal isolated cell types that are preferentially distributed along functional gradients, in large part anchored at each end, the large-scale organization of cortex is likely most apparent when simultaneously considering the spatial distribution of multiple cell types. To visualize this property of cellular organization, we took the difference between Ex1 and PVALB spatial distributions within each parcel, as the former are distributed around transmodal territories and the latter within unimodal regions (Fig. 2B). This combination aligns with the first functional gradient. We further examined the combinatorial alignment of all the cell types to gradients in the next section.

### Multivariate cellular profiles track the functional gradient architecture of cortex

Our initial analyses established spatial relationships between specific cell-types, studied in isolation, and the first two functional gradients of connectivity^22^. However, the extent to which the macroscale functional organization of cortex may be reflected in the spatial distribution of multiple cell types remains to be established. To examine the presence of these multivariate cellular profiles, we elected to use permutational canonical correlation analysis (PermCCA)^50^, which seeks the linear combination of cell type distributions that are maximally correlated to each individual gradient. Inference on the resulting canonical variates was performed using spin-test permutations^42^ on each functional gradient to account for spatial autocorrelation.

Here, the resulting cell type composite scores reflect the linear combination of spatial distributions across all cell-types that are maximally correlated to each functional gradient. As displayed in Fig. 3, we observed cell type composite scores were correlated with both the first and second gradient across both visual and frontal cortex single-cell data (Lake DFC, Fig. 3A,D; Lake VIS, Supplementary Fig. 3A,D). In Fig. 3B,E (for Lake DFC; Lake VIS, Supplementary Fig. 3B,E), we display the strength of involvement of each cell type composite score associated with the first and second functional gradient projected to the cortical surface. At the univariate level, as reported above, a common spatial distribution for these gradient-associated cell types emerged where the profile of enrichment is anchored at one end of the gradient (Fig. 2). Consistent with this pattern, as reflected in our PermCCA cell-type loadings, Gradient 1 (Lake DFC: r=0.524, p_FDR_= 0.004; Lake VIS: r=0.566, p_FDR_=0.002) is most positively correlated with In1, Ex1, and most negatively correlated with PVALB and End; whereas Gradient 2 (Lake DFC: r=0.586, p_FDR_=0.004; Lake VIS: r=0.650, p_FDR_=0.002) most positively correlates with Ex4 and End, and most negatively correlates with Ex3.

**Fig. 3.**
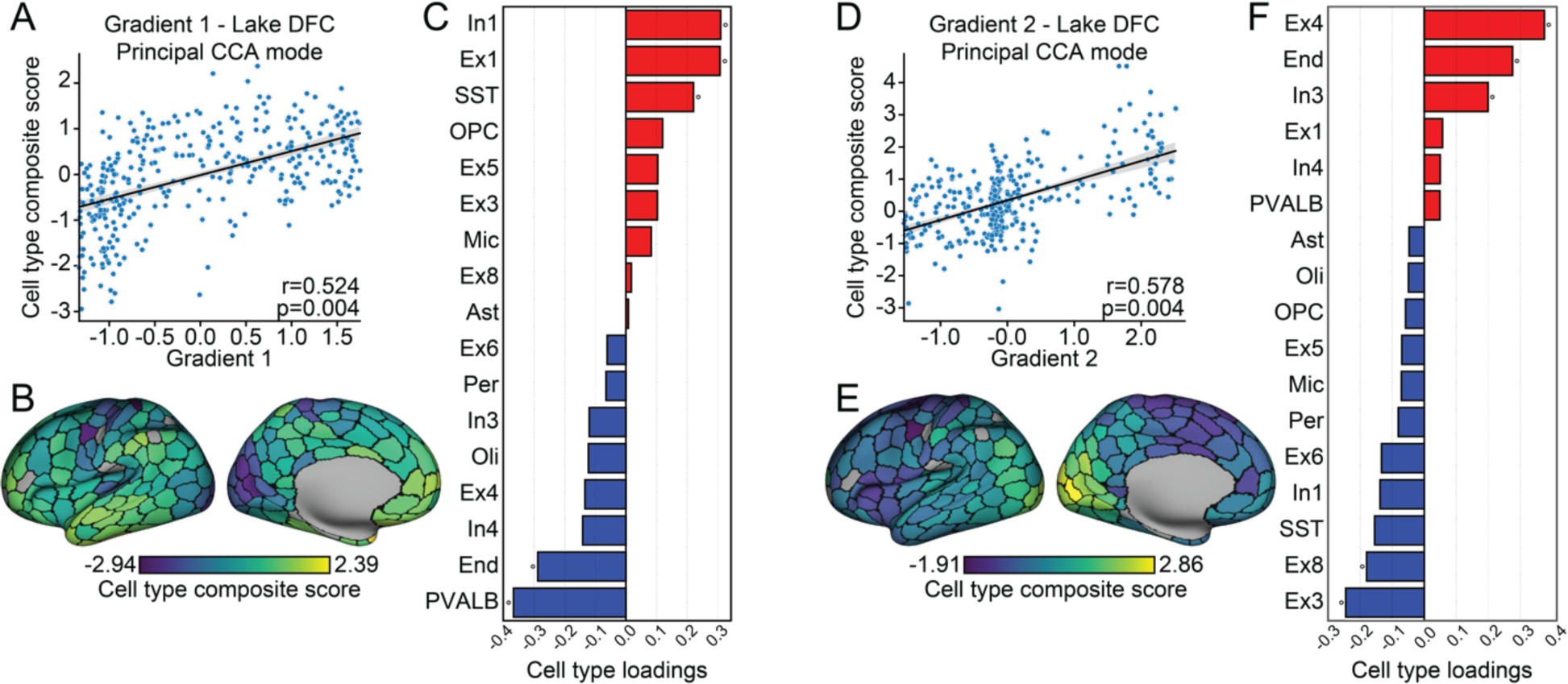
Multivariate cellular profiles follow the macroscale organization of cortex. **A.** The scatter plot displays results of permutational canonical correlation analysis where the first functional gradient was positively associated with a composite score of cell type abundance. **B.** Cell type composite score associated with the first functional gradient projected to the cortical surface. **C.** Canonical loadings of each cell type to the composite score implicate In1, Ex1, PVALB and the SST interneurons and endothelial cells (End; red indicates positive associations; blue, negative associations). **D.** The second functional gradient was positively associated with a cell type composite score of cell type abundance. **E.** Cell type composite score associated with the second functional gradient mapped to the cortical surface. **F.** Canonical loadings of each cell type to the composite score significantly implicated Ex4, Ex3 (marked by gene NEFM) and Ex8 (MCTP2, NR4A2) excitatory neurons, and In3 (TSHZ2, SHISA8) interneuron, and End.

Although the cell types that were primarily associated with gradients in isolation also preferentially contribute the signal to the multivariate analyses, the variance explained with using a composite score now increases to 27.5% (Gradient 1) and 34.3% (Gradient 2; Fig 3A,D) from 9.4% through 14.1% in the single cell-types emerging from the univariate analyses (Fig. 2D). These data suggest that, rather than being specific to isolated classes of cells, the observed topographic similarities may be most apparent when considering the combined spatial profiles of multiple cell types across cortex. To test this idea, we repeated the PermCCA iteratively excluding different combinations of cells that reflect the major contributors for Gradient 1 and Gradient 2 (see Supplementary Table 1-2). Critically, when simultaneously removing all the significant gradient associated cell types revealed in the univariate analysis (Gradient 1: In1, Ex1, PVALB; Gradient 2: Ex4), the PermCCA results held for Gradient 1 (Lake DFC: r=0.419, p_FDR_=0.029; Lake VIS: r=0.538, p_FDR_=0.002) and almost held for Gradient 2 (Lake DFC: r=0.497, p_FDR_=0.060; Lake VIS: r=0.546, p_FDR_=0.003). Therefore, while certain cell types may preferentially follow the gradient architecture of human cortex, the observed spatial relationships are robust and broadly conserved across a host of cell types (see cell type abundance correlation matrix in Supplementary Fig 4).

### The cellular composition of large-scale functional networks

Certain cell-types may possess a preferential relationship with specific functional networks. For instance, we observed the increased presence of PVALB cells in the somato/motor and visual networks (Fig. 2D), confirming a similar pattern in prior work^16^. To explicitly assess these network-cell relationships, we calculated an enrichment score for each cell type (see Methods) across the 7 canonical functional networks^23^ (Fig. 4A). Similar to single-cell gradient analyses above, PVALB, End, and oligodendrocyte (Oli) cell types exhibit heightened enrichment within unimodal somato/motor and visual networks, whereas Ex4 is preferentially enriched in visual networks over the other unimodal territories. Conversely, In1, Ex1, Ex5 (including HS3ST5^+^PCP4^-^ Ex5a and HS3ST5^-^PCP4^+^ Ex5b)^41^, SST, astrocyte (AST) and oligodendrocyte progenitor cells (OPC) are most enriched in default and limbic networks, but also broadly across attention network parcels. Of note, some cell types did not cluster within the unimodal or association regions (default, limbic) that anchor the first gradient. Ex8 (CBLN2^+^NR4A2^+^)^41^ cells, for instance, were preferentially enriched in the ventral attention network.

**Fig. 4.**
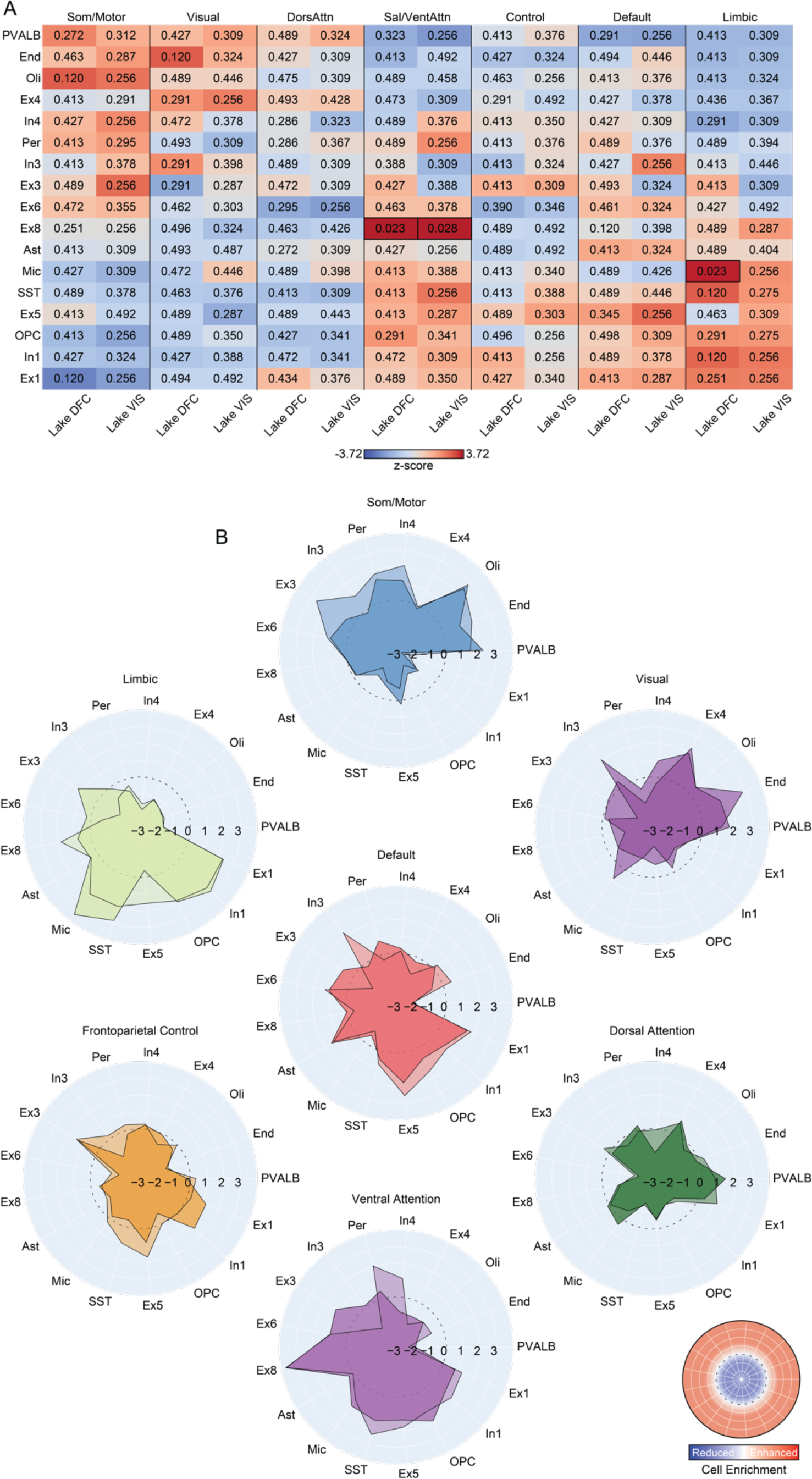
Large-scale functional networks demonstrate distinct cellular profiles. **A.** The table displays the relative cell type enrichment, or absence, within each canonical functional network. Networks are ordered as their estimated position along the first principal gradient^22^. Empirical abundance for each cell type was aggregated within each of the 7 large-scale functional networks. Corresponding null distributions were constructed from parcel-level spin-test, which accounts for spatial autocorrelation^51^. Table fill colors reflect z-scores, derived via subtracting the mean of the null from the observed empirical abundance then dividing the difference by standard deviation of null distribution. Here, z-scores index empirical enrichment relative to the null. Warm colors indicate positive values, numbers in each cell represent the FDR corrected p_spin_ of relative enrichment or absence. Reflecting the presence of a cell-type enrichment gradient spanning between somato/motor and limbic networks, each network shows a unique cell type profile. Marked boxes reflect significant enrichment (p_FDR_<0.05). **B.** Polar plots of z-score across 17 cell types for each network suggest the potential of cellular profiles that may serve as fingerprints that can distinguish each functional network. Score above zero lines (dashed) indicate when a cell type is enriched within a given network relative to the overall distribution across cortical parcels, whereas below zero reflects the relative absence of a cell-type. Polar plot corresponds to imputed cell densities from Lake DFC and Lake VIS are stacked together, with the overlapping area in darker color.

Each column in Fig. 4A shows cellular enrichment profiles across large-scale functional networks. Although the gradient properties of cellular organization are evident from unimodal through association territories, visually distinct network-level enrichment profiles are also apparent, embedded within this sweeping organizational motif. This is further reflected in the polar plots in Fig. 4B, displaying enrichment scores across cell types for each network. Situated at distinct ends of the first functional gradient of connectivity (Fig. 1B), both the unimodal somato/motor and visual networks and the default and limbic networks exhibit the most contrast between heightened or reduced enrichment profiles across cell types, with the ventral attention network also exhibiting pronounced variability in cellular enrichment. Broadly, the presence of distinct cellular fingerprints across functional networks suggests that cellular profiles derived from most-mortem tissue samples may reflect, and be predictive of, the functional allegiance of a given cortical parcel.

### Predicting network allegiance from cell-type abundance in post-mortem tissue

Our findings raise an important question: can cellular profiles imputed from bulk tissue samples be used to directly infer *in vivo* properties of brain organization? The above analyses identify individual cell classes that preferentially follow the spatial topography of large-scale functional networks, suggesting the presence of network specific cellular fingerprints. We use these results as a foundation to directly test the extent to which parcel-level multivariate cellular profiles can be used to predict their corresponding functional network assignments, as derived through fMRI. Support vector machines (SVM) were trained to predict the functional network allegiance of post-mortem tissue samples from parcel-level cell type abundance (see Methods). Performance of models trained from empirical data were compared to a comprehensive set of increasingly stringent null models: (1) theoretical chance of predicting correctly given that the parcel is from certain network (1/7, p_chance_); (2) through models trained from network labels that are randomly permuted (p_perm_) or (3) shuffled while controlling for spatial autocorrelation (p_spin_). Here, we focus our interpretation on the most stringent significance null-model (p_spin_). The network-and parcel-level prediction accuracy and alternate significance thresholds are displayed in Fig. 5.

**Fig. 5.**
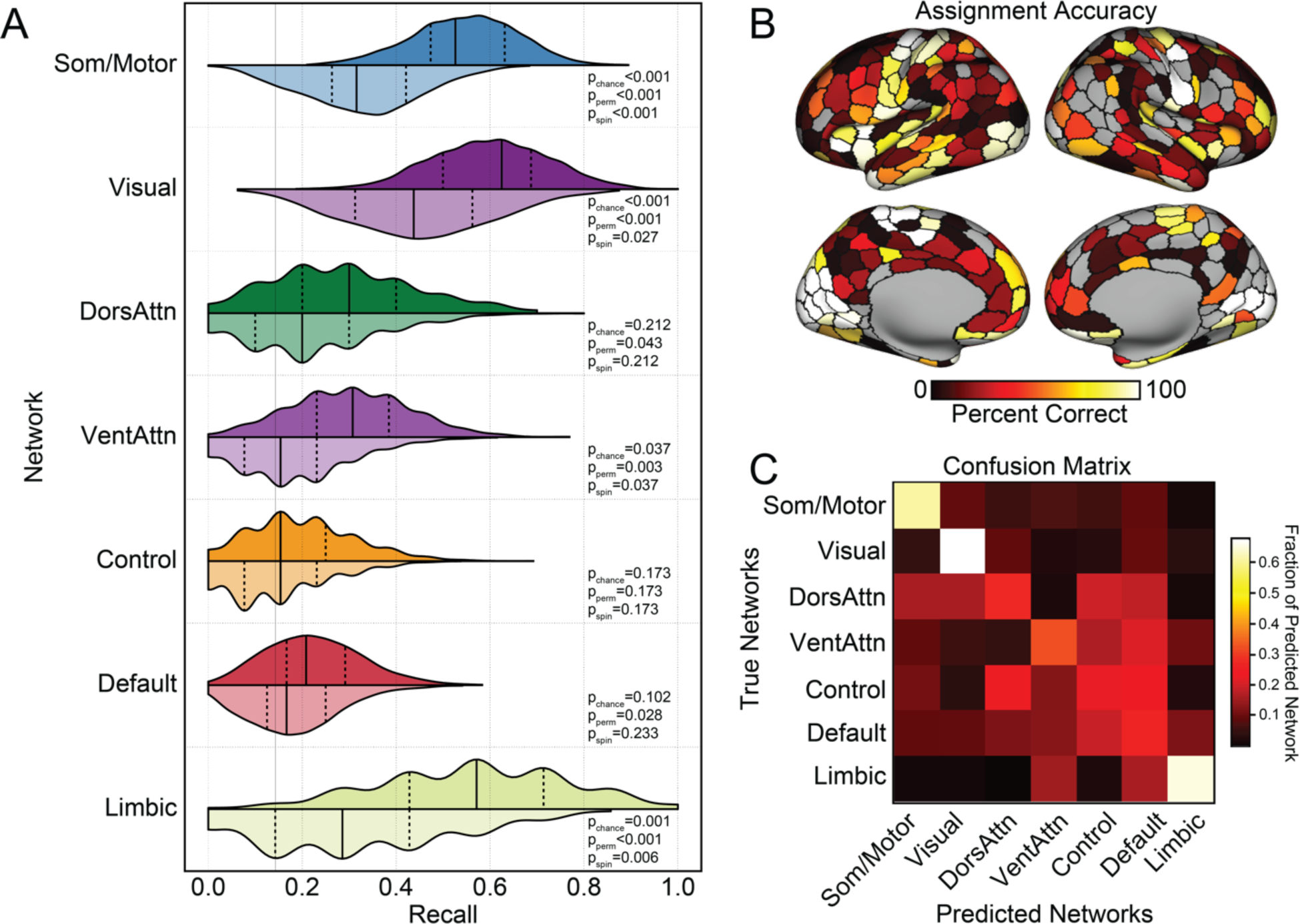
Large-scale functional network assignment can be predicted by cell-type abundance in post-mortem tissue. **A.** Histograms display the SVM recall, or the probability of correctly classifying a parcel to the associated network. These data suggest the classifiers were able to predict somato/motor, visual, ventral attention, and limbic networks significantly above chance. Distributions in darker color were constructed from 1000 classifiers trained on real network labels, and the lighter colored distribution represent classifiers trained on network labels shuffled by spin-test that controls for spatial autocorrelation^50,52^. The solid lines indicate median and the dashed lines represent quartiles of the distribution. Significance of empirical recall were constructed relative to thresholds of increasing stringency, (1) theoretical chance (1/7, p_chance_); (2) models trained from randomly permuted network labels (p_perm_) (3) or labels shuffled by spin-test (p_spin_). **B.** Accuracy of network assignment across cortical parcels, calculated from all testing sets. **C.** Each row of the confusion matrix represents the fraction of parcels within the specific network that were predicted as belonging to each of the 7 networks. The diagonal represents the percentage of correctly classified parcels within each network. Here, the confusion matrix suggests a preferentially distinct cellular profile for somato/motor, visual, and limbic networks. While classification accuracies were low for the remaining association cortex networks, dorsal attention, default, and control networks display a higher rate of misclassifications among each other, relative to unimodal networks.

The SVM model was able to successfully decode parcel-level network assignments across cortex (empirical F1_median_=0.411, null F1_median_=0.252, p_spin_<0.001, distribution plots in Supplementary Fig. 5), indicating that inferred cellular abundance from resected post-mortem tissue reflects functionally relevant properties of brain organization. When considering individual networks, the SVM models trained on parcel-level imputed cell densities successfully predicted somato/motor (p_spin_<0.001), visual (p_spin_=0.027), ventral attention (p_spin_=0.037) and limbic (p_spin_=0.006) networks (all other p_spin_>0.173, Fig. 5A). When projecting parcel-level accuracies to the cortical surface (Fig. 5B), within network variability was evident, indicating spatial heterogeneity of the cellular composition and associated network level assignment accuracies [somato/motor (average parcel accuracies±standard deviation: 0.605±0.390); visual (0.674±0.390); limbic (0.654±0.371); dorsal attention (0.271±0.238); ventral attention (0.323±0.350); default (0.264±0.246); and control (0.225±0.236); detailed in Supplements table2)]. The confusion table, presented in Fig. 5C, highlights the assignment stability of the somato/motor, visual, and limbic networks. Of note, while assignment accuracy is reduced in the three remaining networks, where misassignment occurs in parcels from control, attention, and default networks, they are likely to be assigned to other association cortex networks. For example, the dorsal attention network borders with, and is interdigitated between, unimodal and transmodal areas. However, when misclassifications occur, dorsal attention parcels are more likely to be labeled as default or control than somato/motor or visual. The SVM results emerging from different data exclusion criteria are displayed in Supplementary Figs 6-10. Together, these data confirm the presence of distinct cellular fingerprints within some functional systems and support the need for additional research into the cytoarchitectonic determinants of network topography.

## Discussion

Since Theodor Hermann Meynert first observed regional variations in the histological structure of gray matter across the cerebral cortex^2^, the resection and study of post-mortem tissue samples has revealed core insights into the cellular composition of the central nervous system. Over the past few decades, methodological advances have made it possible to map the macroscale organization of brain functions *in vivo*, providing the potential for deep biological insight into the genetic, molecular, and cellular bases of cortical brain organization. Here, integrating transcriptional and neuroimaging data, we demonstrate that imputed cell type distributions follow the hierarchical functional architecture of the cortical sheet. Select cell types were found to spatially couple with aspects of cortical gradient and network organization, which was most evident in a distinction between higher-order association and unimodal territories. Suggesting that regional variation in cellular profiles may reflect the layered aspects of cortical organization, multivariate cellular fingerprints captured a substantial portion of the spatial variability in both functional gradient topographies and parcel-level network assignments. Finally, imputed cell type densities, derived from post-mortem tissue samples, could be used to accurately predict parcel-level network assignments, suggesting the presence of cellular markers of network-level brain functions as assessed through rs-fMRI. Together, these results indicate a close link between the functional organization of cortex and spatial variability of cell-type distributions with important implications for the study of the cellular basis of brain functions across health and disease.

Recent work suggests that the distributions of select cell gene markers may spatially couple to regional differences in functional MRI signal variability. For instance, single-marker and polygenic cell deconvolution has established a spatially dependent relationship between heritable variance in *in-vivo* functional MRI signal amplitude and the topography of parvalbumin expression in post-mortem brain tissue^16^. Here, we extend upon this work, demonstrating the presence of spatial alignment between regional cell densities, imputed from post-mortem tissue, and the functional gradient architecture of cortex. Inhibitory neuron In1, including subtypes as CCK, RELN and VIP, preferentially align with the association end of the principal functional gradient (Gradient 1), whereas PVALB are generally enriched within unimodal areas. The presence of a dichotomous relationship between these two groups of interneurons, embedded within a hierarchical somato/motor-association gradient in adults, echoes their positioning during early embryonic development. The properties of interneuron subtypes are, in part, determined by their spatial origin in the embryonic ganglionic eminence^53^. In rodents, interneuron destined cell types originate from distinct embryonic progenitor zones in the ventral telencephalon. Neural progenitor cells in the medial ganglionic eminence (MGE) give rise to PVALB, whereas progenitors in the caudal ganglionic eminence (CGE) generate RELN and VIP^54^. After neurogenesis, these two broad cell classes are governed by separate transcriptional cascades that direct their tangential migration, layer-specific positioning, and maturation^55,56^. For example, the MGE-originated PVALB is sequentially regulated by transcriptional factors *Nkx2.1*, *Lhx6*, and *Sox6*. As a vastly different regulator for the other interneuron groups, *Prox1* plays a crucial role in allocating the CGE-derived VIP and RELN into superficial layers, and in regulating the subsequent circuit integration and refinement. In addition, excitatory neuron Ex1 also aligns with the association endpoint of Gradient 1. Its marker, *CBLN2*, whose prenatal upregulation coincides with synapse formation onset, promotes dendritic spine formation^47^. Consistent with preferential expansion of prefrontal regions across our evolutionary lineage^57^, the enrichment of the CBLN2 in prefrontal cortex is increased in human primates, relative to rhesus macaques^58^.

Indicating the presence of multivariate cellular profiles that follow the macroscale functional organization of cortex, PermCCA analyses revealed spatial correspondence robustly linking functional gradient architectures with imputed cell-type densities. When considering Gradient 1, the topography of cell type distributions is aligned with the primary unimodal-transmodal gradient of connectivity, reflecting 27.5% of the variance in functional connectivity. Intriguingly, as alluded to above, the spatial patterns of cellular enrichment in adult post-mortem tissue samples seems to mirror the relative spatial distribution of these cell types within their respective embryonic progenitor zones. Interneuron cells enriched in the association areas in adulthood, as one example, include In1, SST, and PVALB. Here, In1 (CCK, VIP, RELN) cells originate within the CGE, SST are derived from the dorsal MGE; whereas the unimodal enriched PVALB are generated from ventral MGE^59,60^. Highlighting the intertwined nature of cellular interactions across cortex, the co-enrichment of excitatory Ex1 (CBLN2^+^) and inhibitory In1 (CCK^+^VIP^+^RELN^+^) in transmodal association areas may reflect a regulatory role of In1 on Ex1 functions. In mice, *CBLN2* serves as an essential extracellular scaffolding protein for VIP-specific inhibitory synaptogenesis^61,62^. These data hint that the cellular basis of the brain’s functional architecture is likely instantiated early in fetal development, providing the opportunity for the study of how patterns of cell migration and maturation shape the cortical functional connectome.

CCK and PVALB inhibitory interneurons differ across a host of morphological and firing properties, forming distinct local inhibitory circuits that can differentially bias network oscillations^63^. Our present analyses highlight a relationship between the primary gradient (Gradient 1) and the relative presence of PVALB and In1 (CCK, VIP, RELN) cells. The distinct computational properties of unimodal- and heteromodal association territories are theorized to reflect the relative preponderance of associated cell types and their specialized roles in primary sensation and cognition^64^. For example, PVALB expressing inhibitory interneurons are preferentially situated in the unimodal visual and somato/motor cortex. PVALB cells are a class of inhibitory interneurons that broadly synapse on the perisomatic region of cortical projection neurons to regulate output^65^. Computational models in rodents suggest that a relative increase in PVALB, relative to SST, may result in stronger feedback inhibition on excitatory neurons within a given patch of cortex. This profile of excitability is thought to allow for short activation timescales that may be optimally suited for processing constantly changing sensorimotor stimuli^16,64,66^. Conversely, enriched in the transmodal association area, CCK interneurons display a broad range of spiking patterns, varying from synchronous transmission, which may enhance precise inhibition timing, through asynchronous repetitive activations that are thought to modulate inhibition strength and signal durations^67–69^. This wide spiking spectrum, well suited for association cortex, enables them to integrate signals from various sources. Although future computational modeling work is warranted, the present data suggest a link between the migration and maturation of local cellular circuits across cortex and the subsequent development and refinement of macro-scale functional systems. Here we show cell-types that link to Gradient 1 and Gradient 2 in adults. Although these cell classes reach their cortical destinations in early life, the associated developmental and maturational trajectories extend from childhood to adolescence^38^. It remains to be determined how cell maturation, refinement, and synaptic connectivity links to the development of functional gradient architectures across the lifespan.

The large-scale network architecture of the cortex includes abrupt transitions that are embedded along continuous functional gradients^22^. Broadly, similar patterns are evident in cortical atlases defined through cell staining and morphological analysis, where homogeneous cell components are evident within local patches while relatively abrupt transitions can occur between some adjacent territories^6^. Evidence has emerged suggesting links between functional network parcellations of cortex and the presence of some cytoarchitectonic boundaries^23^. Here, functional network defined boundaries and parcels have been observed to adhere to select histologically defined areas, including Broca’s area (area 44) and aspects of the postcentral sulcus (area 2 and 3)^35^. In the present analyses, using the imputed abundance across neuronal and nonneuronal cells in post-mortem tissue, we demonstrate that the functional network allegiance across cortical parcels can be generally predicted. Although parcel-level prediction was broadly evident across cortex, substantial spatial heterogeneity was evident in the relations linking cellular profiles with network assignments. The somato/motor, visual, limbic and ventral attention networks exhibited the most distinct and predictable cellular profiles. In contrast, control, dorsal attention and default networks displayed smoother and/or more heterogeneous spatial transitions across cell types. Underscoring the translational potential of multi-scale neuroscience approaches, these findings bridge levels to demonstrate predictive relationships connecting cellular profiles across the cortex with the *in vivo* functional organization of the human brain.

The present work should be interpreted in light of several limitations. First, the reported cell type abundance are imputed from the bulk tissue microarray data based on the gene expression signature constructed from single-nucleus RNA sequencing. The microarray approach does not provide direct estimates of gene transcription, rather, here we examine within-probe differences across samples. To obtain estimates robust across subjects, bulk samples require aggregation into parcels, limiting the spatial resolution. As parcellation and cell-type definitions improve the pattern observed will likely unfold in more detail. Second, to control for spatial heterogeneity of single-cell signature, only the cell types common on both Lake DFC and Lake VIS are studied here. This may have resulted in an underestimation of the true relationships linking the spatial distributions of cell types and brain functions. In mice, interneuron cell types are broadly conserved across cortical regions, while pyramidal cell diversity shows higher spatial variability^70^. The reported imputed spatial distribution of cell types common in both single-cell samples (Lake DFC and Lake VIS) showed robust patterns, but it remains unclear how the cell type diversity varies across the human cortex. As single-cell samples covering more cortical regions are gradually developed, future work should incorporate these spatially variable profiles when considering cell type abundances. Finally, in the present work cells are defined from transcriptomics. As cells may be defined though their transcriptional profiles, morphology, or firing patterns^65^, follow-up studies should consider how to best integrate these diverse cell definitions in analyses. Here, researchers should consider alternate organizational models of brain functioning (e.g., additional network atlases, graph theoretical models of cortical information processing, anatomical and diffusion defined network solutions, etc).

The present results demonstrate that the functional gradients and networks of the cerebral cortex are linked to spatial variability in cellular profiles. These data suggest that the imputed cell type densities from post-mortem tissue capture global patterns functional connectivity as assessed through rs-fMRI, revealing the potential to bridge across *in vivo* and *ex vivo* methods in the study of human brain functions. Collectively, these findings highlight a connection between the functional organization of the cortex and its cellular underpinnings, which has significant implications for understanding the cellular basis of brain functions in both healthy and diseased states.

## Methods

### Functional Connectivity Gradient Analysis

Gradients, or components with similar functional connectivity patterns, were derived in a manner consistent with Margulies et al.^22^. Briefly, functional connectivity matrices averaged from 820 subjects in HCP dataset^39^ coregistered via MSMAll were downloaded from ConnectomeDB^71^(https://www.humanconnectome.org/storage/app/media/documentation/s900/8 20_Group-average_rfMRI_Connectivity_December2015.pdf). The 10,242 x 10,242 per hemisphere cortical FC matrices (represented in fsaverage5 surface space^72^) for each subject was calculated from 1-hour resting-state fMRI concatenated from four minimally preprocessed^73–76^, spatially normalized 15-min scans. From these group-averaged FC matrices, correlation coefficients were Fisher z-transformed via a hyperbolic tangent function to scale the value between -1 and 1. The top 10% connections of each vertex were preserved and all other values were set to 0 to enforce sparsity. The cosine distance between any two rows of the FCz matrix were calculated then subtracted from 1 to generate a symmetrical similarity matrix. Gradients were derived from the similarity matrix by diffusion map embedding, as validated and detailed in^22,38^(https://github.com/satra/mapalign). This approach nonlinearly projects high-dimensional functional connectivity into a low-dimensional space. Here, a gradient reflects an axis of FC variance along which cortical vertices fall in a spatially continuous order, with adjacent vertices sharing similar geographically short- and long-range correlations to the rest of the cortex. The two gradients explaining highest variance were selected for subsequent analysis. Vertex-level gradients were averaged across the 400 cortical parcels in the Schaefer functional atlas^35^.

### Functional Parcellation Analysis

To characterize the functional network structure of the cortical sheet, we used 400 roughly symmetric ROIs from 7 specific brain networks^23^ in the left and right hemispheres as derived through the cortical parcellation of Schaefer and colleagues^35^. The functional networks used here were previously derived and validated using data from 1000 adults in the Genomics Superstruct Project (GSP)^77^ as detailed in^23,35^. In short, each network is a cluster of vertices that shares homogenous resting-state fMRI FC to the rest of the cortex.

### Brain Gene Expression Data Processing

Human microarray gene expression data obtained from bulk samples of 6 postmortem brains were downloaded from AHBA dataset (http://human.brain-map.org/)^11^. Raw data were processed using the abagen toolbox (https://github.com/netneurolab/abagen)^78,79^ at the sample level, following the practice recommended by Arnatkevičiūtė et al^80^ and implemented by others^13^. Probes were reannotated using data provided by Arnatkevičiūtė et al^78^, and those without Entrez IDs were excluded. Probes that exceed background noise in 30% of all tissue samples were included, among which the probe with highest differential stability for each gene was selected. 16,383 genes were retained after the processing. For each donor, tissue sample expression values were first z-scored across genes and then these gene expression values were then z-scored across samples. Consistent with Anderson et al^16^, individual cortical tissue samples were mapped to each AHBA donor’s Freesurfer derived cortical surfaces, downloaded from Romero-Garcia and colleagues^81^. Native space mid-thickness surfaces were transformed to a common fsLR32k group space while maintaining the native cortical geometry of each individual donor. The native voxel coordinate of each tissue sample was mapped to the closest surface vertex using tools from the HCP workbench. Tissue samples were included if they were collected from less than 4 mm from the nearest surface vertex, resulting in 1,676 analyzable cortical samples.

### Cell Type Deconvolution

Cortical cell type abundance distributions were inferred following the procedures detailed in Anderson et al^16^. In brief, single-nucleus droplet-based sequencing data obtained by Lake et al. were downloaded from Gene Expression Omnibus website (“GSE97930” [https://www.ncbi.nlm.nih.gov/geo])^41^. Count matrices derived from unique molecular identifier (UMI) were preprocessed via Seurat^82^, where outlier cells and minimally expressed genes were filtered then the data were log-normalized. Genes were referred to by Entrez IDs, among which only the IDs shared by both Lake and AHBA datasets were included. The superordinate cell identities defined by Lake et al^41^ were applied for categorizing transcriptionally similar cell types, to reduce the collinearity. After processing, the snDrop-seq data were de-log-transformed before feeding into CIBERSORTx(https://cibersortx.stanford.edu/)^83^ as reference for cell type abundance imputation on each AHBA bulk tissue sample. Gene signature matrices for 18 cell types were derived from visual (Lake VIS) and frontal (Lake DFC) samples separately. Cell type abundances were consequently imputed from across AHBA samples taking each signature matrix as reference per donor. The correlation between Lake VIS and Lake DFC derived gene signatures was validated in Anderson et al^16^.To further minimize the effects of spatial heterogeneity of single-cell transcriptional signatures, later analysis only utilized common cell types between Lake DFC (excluding Ex2) and Lake VIS (excluding In2), which is in total 17 cell types. The cell type abundance for each AHBA cortical sample were mapped to the cortical vertices represented in fsaverage6 surface space then parceled into the Schaefer 400 atlas^35^, first at the individual level and then averaged across donors.

### Identification of Cell Types Spatially Correlated to Functional Gradients

The spatial pattern of each of the 17 cell type abundances parceled in the Schaefer atlas was correlated (Spearman’s Rho) with the primary and secondary functional gradients parceled using the same atlas. The statistical significance of correlation was assessed through spin-test^42^, which permuted the gradient at vertex level (represented in fsaverage5) for 1000 times while reserving the spatial auto-correlation. The permuted gradient and cell-type correlations were used to construct a null distribution of correlation values. The cell type abundances inferred based on the visual and frontal single-nucleus samples were tested separately and were FDR corrected for 34 multiple comparisons (17 cell types x 2 gradients).

### Examination of Cell Types Combinational Correlation to Functional Gradients

Permutation canonical correlation analysis (PermCCA) was used to investigate the multivariate relationship between spatial distribution of cell types and each gradient. CCA allows us to examine the linear combination of all the cell type abundances that maximally correlate with each gradient. The statistical significance of the canonical variates were tested via a permutation method that controls for cortical spatial autocorrelation (https://github.com/andersonwinkler/PermCCA/tree/master)^50^, where the null distributions of gradients were generated from spin-test described in the previous section. Then FDR was applied to correct for multiple comparisons (2 gradients). The cell types’ linear combinational correlation to the first two gradients was measured by cell type composite score. Each cell type’s contribution to this correlation was measured by loadings, the correlation between cell type abundance distribution and gradient canonical variate.

PermCCA was repeated with removal of different combinations of the cell types that were each univariately correlated with Gradient 1 and 2. Each group of PermCCA results were FDR corrected for the number of removal combinations (12 for Gradient 1 with Lake DFC imputed cell types, 10 for Gradient 1 with Lake VIS imputed cell types; 3 for Gradient 2 with Lake DFC imputed cell types, 1 for Gradient 2 with Lake VIS imputed cell types).

### Cell Types Enrichment in Functional Networks

Each one of the 400 cortical parcels was assigned to a functional network in a validated 7-network solution derived by Thomas Yeo^23^. Across each cell type, the empirical abundances were averaged across parcels within a given functional network. The empirical abundances were then permuted across the cortex for 1000 times, controlling for spatial autocorrelation, yielding 1000 null models for each cell type. Given that the cell type abundances were aggregated at parcel level, the Cornblath version of spin-test^51^ was used as recommended by Markello and Misic^84^, which projects parcel abundances to vertices, rotates, and takes the mean of vertices in each parcel. The same within-network cell type abundance averaging process was repeated on these null models, generating a null distribution of mean abundance for each cell type within each network. For each type of cell, the enrichment score of a network was calculated by taking the difference between the empirical abundance and the mean of null abundance distribution, then divided by the standard deviation of the null abundance distribution. P-values were first calculated from two-tailed tests then FDR corrected for 119 multiple corrections (17 cell types x 7 networks).

### Cell Types Predicting Functional Networks

Support vector machines (https://scikit-learn.org/stable/modules/svm.html#svm-classification) were trained to predict the functional network each cortical parcel belongs to based on abundances of 17 cell types within that parcel. Since two out of six donors in AHBA had samples from the right hemisphere, models were trained on parcels from only the left hemisphere and from both hemispheres as two parallel groups. Within each group, three sub-groups of models were trained separately, based on the cell type abundance imputed from independent single-cell samples, Lake DFC and Lake VIS, as two replicates. Two sub-groups were trained from the two replicates respectively and one sub-group ensembled the information from the two replicates. For each model, parcels were randomly shuffled and split into 1000 distinct train (70%) and test (30%) sets without replacement. Given that the number of parcels within each functional network are not balanced, the train-test split was stratified within each network category. Nested three-fold cross-validation was implemented to select and validate the hyperparameters in the training set. Kernels and regularization parameters were first selected and tuned in the inner 2-fold cross-validation, the models’ performance were then evaluated in the outer 3-fold cross-validation, the final model was the parameter combination with the highest score. The regularization parameter was set to adjust weights inversely proportional to class frequencies in the training data to control for the unbalanced class size within each split. F-1 score was used to evaluate the models’ overall performance. Recall, which indexes the probability of correctly classifying a parcel given it is from a certain class, was used to evaluate the predicting performance within each class. Accuracy for each parcel, calculated by total number of times it was correctly classified over total number of the times it was included in the test set. These metrics were evaluated from the 1000 test sets.

The predictive metrics for every model were evaluated against models fitted from permuted network labels^85,86^. A Hungarian version of spin-test was applied, which uniquely reassigns each parcel’s network label for every rotation that controls spatial autocorrelation^52,84^. Each permutation was used to train and test a null model using a randomly selected hyperparameter combination from the set of 1000 optimal hyperparameter combinations for the original model^87^. Prediction performance metrics from each of the original model’s 1000 train-test splits were then compared to the median prediction accuracy from the null distribution. Consistent with prior works^85,86^, the p-value for each metric’s significance is defined as the proportion of 1000 original models with performance score less than or equal to the median performance of the null model. Performance metrics were considered to be significant if they performed better than the median null performance for more than 950 of the 1000 original models.

Code and associated cell type abundance maps will be publicly available upon publication.

## Author contributions

X.H.Z. and A.J.H. designed the research. X.H.Z. conducted the research. X.H.Z., K.M.A., H.M.D., S.C., E.D., P.E., D.M., and A.J.H. analyzed and interpreted the results. X.H.Z. and A.J.H. wrote the paper, which all authors commented on and edited. X.H.Z. and A.J.H. made figures. X.H.Z. analyzed the data. X.H.Z. published the code. All authors provided analytic support.

## Acknowledgements

This work was supported by the National Institute of Mental Health (Grants R01MH120080 and R01MH123245 to A.J.H.). This work was also supported by the following awards to E.D.: the Yale University Kavli Institute for Neuroscience Postdoctoral Fellowship for Academic Diversity, the Northwell Health/Feinstein Institutes for Medical Research Advancing Women in Science and Medicine Career Development Award and the Feinstein Institutes for Medical Research Barbara Zucker Emerging Scientist Award. Analyses were made possible by the high-performance computing facilities provided through the Yale Center for Research Computing. This work used data from the Allen Institute for Brain Science. Additional data were provided, in part, by the Human Connectome Project, WU-Minn Consortium (Principal Investigators: David Van Essen and Kamil Ugurbil; 1U54MH091657) funded by the 16 NIH Institutes and Centers that support the NIH Blueprint for Neuroscience Research; and by the McDonnell Center for Systems Neuroscience at Washington University. This manuscript reflects the views of the authors and may not reflect the opinions or views of the Allen Institute, NIH, or the HCP consortia investigators.

---

**Supplementary Table 1.** Multivariate cellular correlation with Gradient 1 after iteratively removing cell types individually associated with Gradient 1. Cell types that are univariately correlated with the first principal gradient are removed from CCA in different combinations. The top row is the CCA performed without the combination of all the Gradient 1-related cell types surviving from significance tests in both Lake DFC and Lake VIS imputed cell type abundances. The rows coming after are CCA performed excluding different combinations of Gradient 1-related cell types surviving from univariate significance tests in Lake DFC (Ex1, In1, End, PVALB) or Lake VIS (Ex1, In1, Ex5, PVALB) imputed cell type abundances. The correlations between multivariate cell types and Gradient 1 remain above 0.419 for all removal combinations, with one exception (removing all significant univariate cell types in Lake DFC: Ex1, In1, End, PVALB). These data suggest the cellular correlation with Gradient 1 is preserved across a host of different cell types. The r values reflect Pearson Correlations. P_FDR_ reflects the p-values emerging from the spin tests following multiple comparison correction with Lake DFC or Lake Vis.

| Multivariate Cell Type Abundances Correlation with Gradient 1 |  |  |  |  |  |
| --- | --- | --- | --- | --- | --- |
| Frontal Cortex Cell Signature (Lake DFC) |  |  | Visual Cortex Cell Signature (Lake VIS) |  |  |
| Removed Cell Types | r | p<sub>FDR</sub> | Removed Cell Types | r | p<sub>FDR</sub> |
| Ex1, In1, PVALB | 0.419 | 0.029 | Ex1, In1, PVALB | 0.538 | 0.002 |
| Ex1 | 0.508 | 0.003 | Ex1 | 0.561 | 0.002 |
| In1 | 0.502 | 0.003 | In1 | 0.548 | 0.002 |
| End | 0.513 | 0.003 | Ex5 | 0.535 | 0.002 |
| PVALB | 0.505 | 0.003 | PVALB | 0.564 | 0.002 |
| Ex1, End | 0.493 | 0.003 | Ex5, PVALB | 0.529 | 0.002 |
| In1, PVALB | 0.477 | 0.008 | Ex1, PVALB | 0.560 | 0.002 |
| Ex1, PVALB | 0.469 | 0.008 | In1, PVALB | 0.546 | 0.002 |
| In1, End | 0.491 | 0.003 | Ex5, Ex1, In1 | 0.519 | 0.002 |
| Ex1, In1 | 0.479 | 0.010 | Ex5, Ex1, In1, PVALB | 0.509 | 0.002 |
| End, PVALB | 0.484 | 0.003 |  |  |  |
| Ex1, In1, End, PVALB | 0.366 | 0.096 |  |  |  |

**Supplementary Table 2.** Multivariate cellular correlation with Gradient 2 after iteratively removing cell types individually associated with Gradient 2. Cell types that are univariately correlated with the second principal gradient are removed from CCA. The top row is the CCA performed without the combination of all the Gradient 2-related cell types surviving from significant tests in both Lake DFC and Lake VIS imputed cell type abundances (Ex4 only). The rows coming after are CCA performed without different combinations of Gradient 2-related cell types surviving from significance tests in Lake DFC (Ex4 and End) or Lake VIS (Ex4 only) imputed cell type abundances. The correlations between multivariate cell types and Gradient 2 remain above 0.497 for all combinations, with one exception (removing all significant univariate cell types in Lake DFC: Ex4 and End).

| Multivariate Cell Type Abundances Correlation with Gradient 2 |  |  |  |  |  |
| --- | --- | --- | --- | --- | --- |
| Frontal Cortex Cell Signature (Lake DFC) |  |  | Visual Cortex Cell Signature (Lake VIS) |  |  |
| Removed Cell Types | r | p<sub>FDR</sub> | Removed Cell Types | r | p<sub>FDR</sub> |
| Ex4 | 0.497 | 0.060 | Ex4 | 0.546 | 0.003 |
| End | 0.509 | 0.060 |  |  |  |
| Ex4, End | 0.393 | 0.220 |  |  |  |

**Supplementary Table 3.** Summary statistics for parcel-level classification accuracy across the seven functional networks. Parcel-level accuracy was obtained from SVM classifiers trained and tested on the samples from both hemispheres, ensembling Lake DFC and Lake VIS imputed cell type abundance. Statistics were calculated from prediction accuracy of parcels belonging to the same network.

| Networks | Mean | Max | Min | Std |
| --- | --- | --- | --- | --- |
| Som/Motor | 0.605 | 1.000 | 0.000 | 0.390 |
| Visual | 0.674 | 1.000 | 0.000 | 0.390 |
| DorsAttn | 0.271 | 0.789 | 0.007 | 0.238 |
| Sal/VentAttn | 0.323 | 0.991 | 0.000 | 0.351 |
| Limbic | 0.654 | 1.000 | 0.000 | 0.371 |
| Control | 0.225 | 0.819 | 0.000 | 0.236 |
| Default | 0.264 | 0.864 | 0.000 | 0.246 |

**Supplementary Fig 1.**
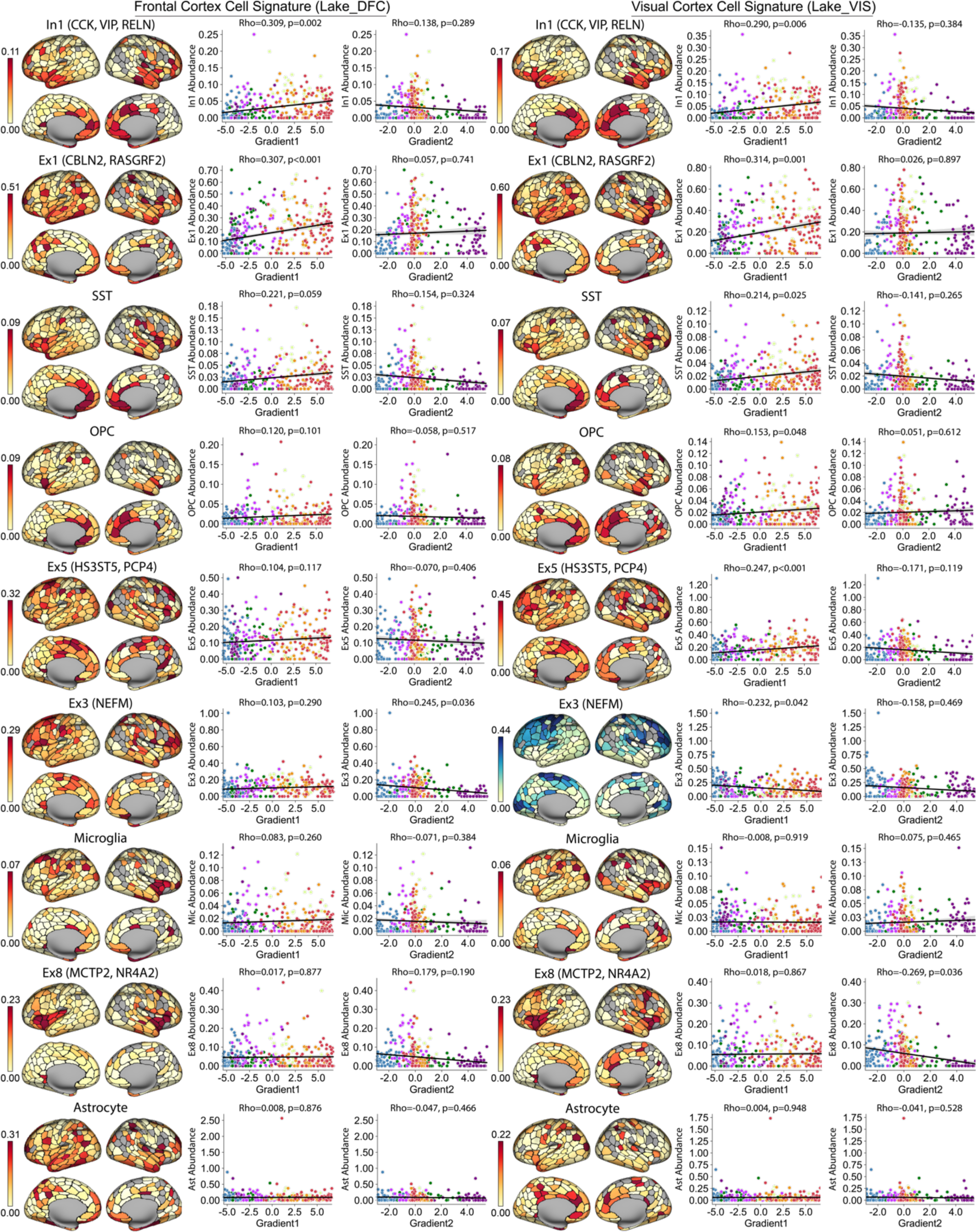
Spatial distribution of cell types preferentially distributed in association cortex. The imputed cell type abundance across cortex in both hemispheres are aggregated in 400 Schaffer parcels. Cell types imputed from single-nucleus droplet (snDrop) samples in the frontal lobe (Lake DFC) are on the left panel and the right is imputed from visual cortex-sourced snDrop samples (Lake VIS). Within each panel, the scatter plot on the left shows the correlation between cell abundance and the first principle gradient (Gradient 1) across cortical parcels, and the correlation with the second gradient (Gradient 2) is displayed on the right. Dots are color-coded by the functional network each parcel belongs to. A positive correlation with Gradient 1 indicates the cell is preferentially spaced on the transmodal association cortex. Such distribution preference reflected on the scatter plot of Gradient 2 is a peak near 0. Though only the top two cells (Ex1 and In1) have significant positive relationships with Gradient 1, the patterns are visually identifiable in the rest of the top five cells (SST, OPC, and Ex5).

**Supplementary Fig 2.**
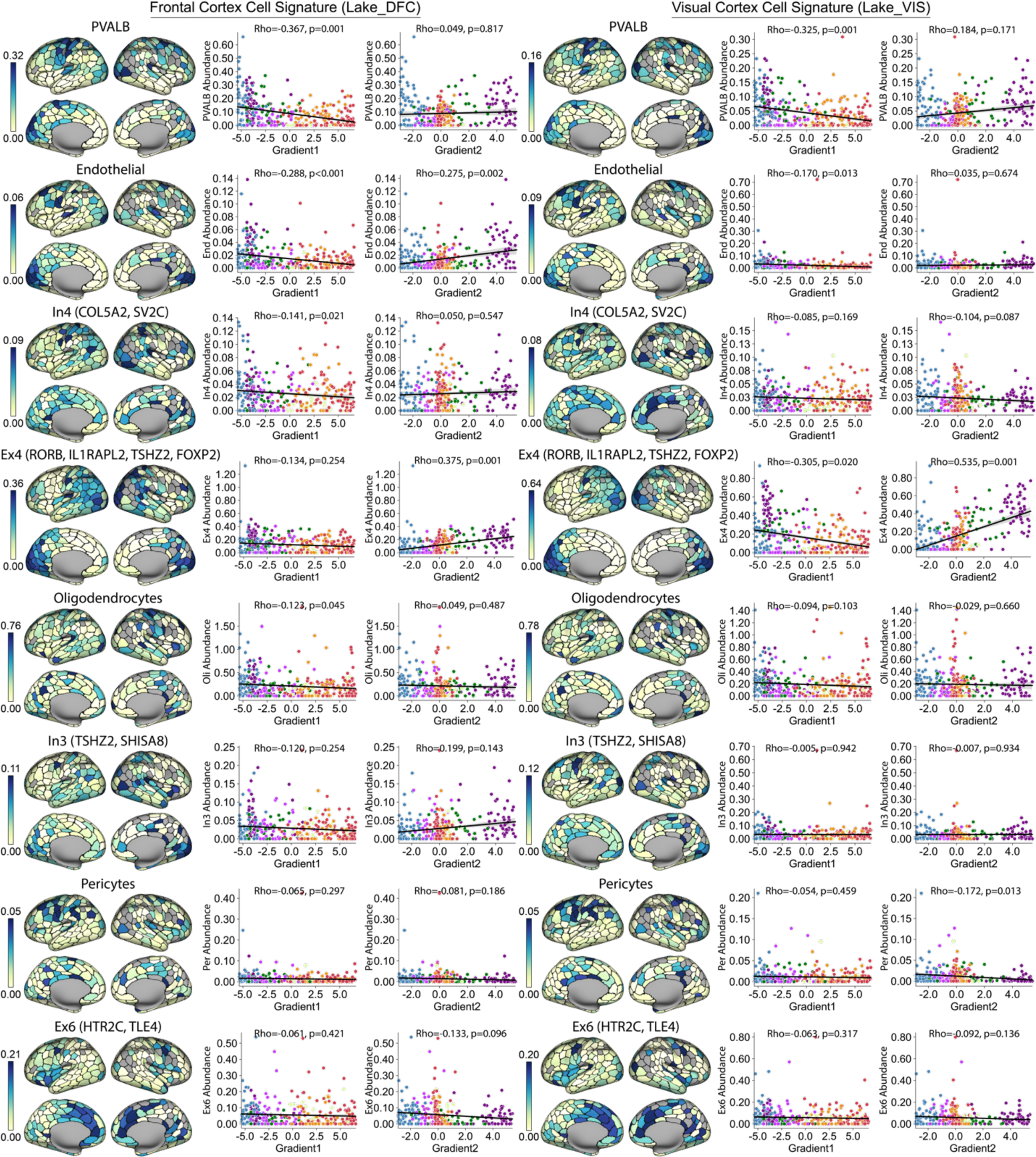
Spatial distribution of cell types preferentially distributed in the unimodal area. The imputed cell type abundance across cortex in both hemispheres are aggregated in 400 Schaffer parcels. Cell types imputed from single-nucleus droplet (snDrop) samples in the frontal lobe (Lake DFC) are on the left panel and the right is imputed from visual cortex-sourced snDrop samples (Lake VIS). Within each panel, the scatter plot on the left shows the correlation between cell abundance and the first principle gradient (Gradient 1) across cortical parcels, and the correlation with the second gradient (Gradient 2) is displayed on the right. Dots are color-coded by the functional network each parcel belongs to. A negative correlation with Gradient 1 indicates the cell is preferentially spaced on the unimodal area. On the scatter plots, this is the peak near the negative end of Gradient 1. On Gradient 2, this is the peak on the visual or somato/motor or both ends. Though only the top two cells (PVALB and End) have significant negative relationships with Gradient 1, the patterns are visually evident in the rest of the top four cells (In4 and Ex4).

**Supplementary Fig 3.**
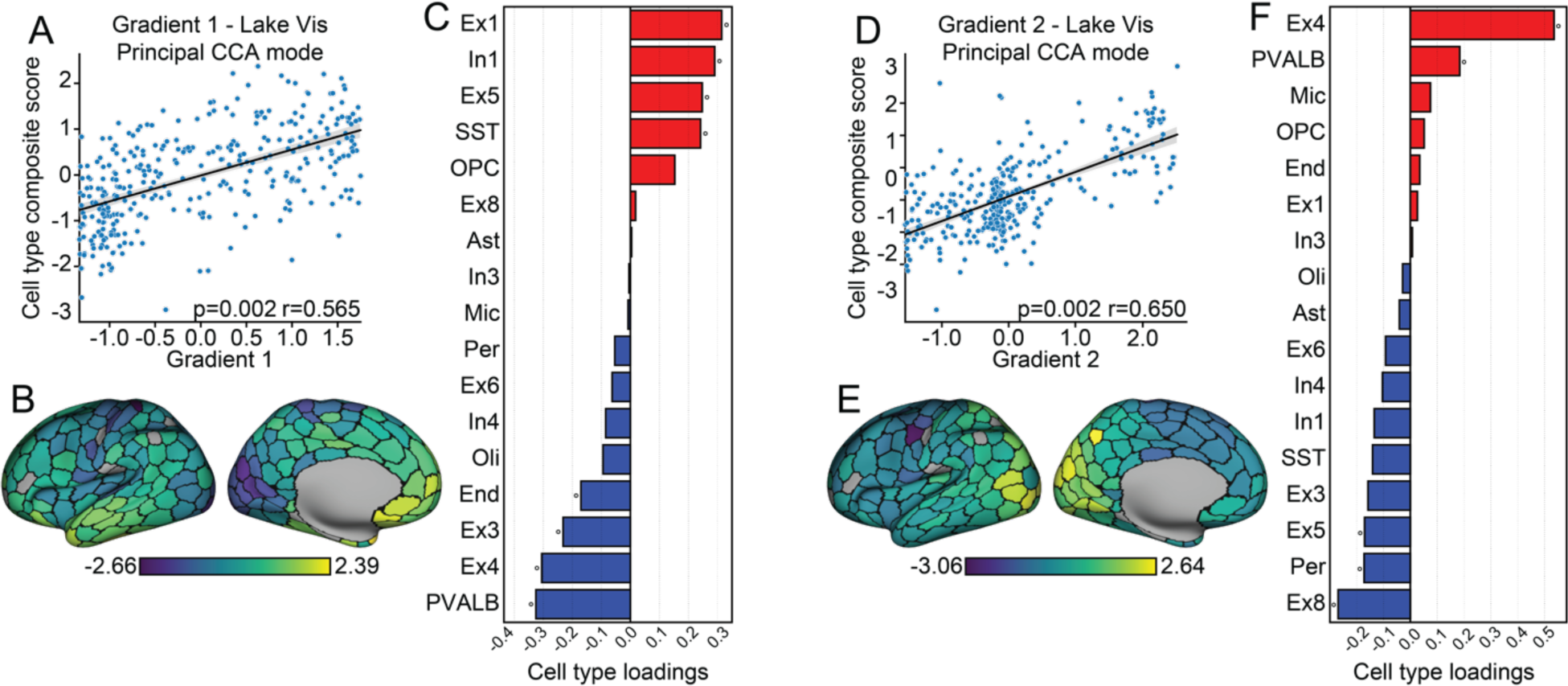
Lake VIS imputed cell type abundance: Multivariate cellular profiles follow the macroscale organization of cortex. The analysis performed based on the Lake VIS imputed cell type abundance showed similar correlation strength and pattern with functional gradients, consistent with the results of Lake DFC (in Fig. 3). **A.** The scatter plot displays permutational canonical correlation analysis modeling results. Across cortical parcels, the first functional gradient was positively associated with the cell type composite score. **B.** Cell type composite score associated with the first functional gradient projected to the cortical surface. **C.** Cell type loadings from the first gradient indicate the correlation is strongest within previously identified Ex1, In1, and PVALB (red indicates positive associations; blue, negative associations). **D.** The second functional gradient was positively associated with the cell type composite score. **E.** Cell type composite score associated with the second functional gradient mapped to the cortical surface. **F.** Ex4 and Ex8 (MCTP2, NR4A2) excitatory neurons remain strong contributors to the correlation with Gradient 2.

**Supplementary Fig 4.**
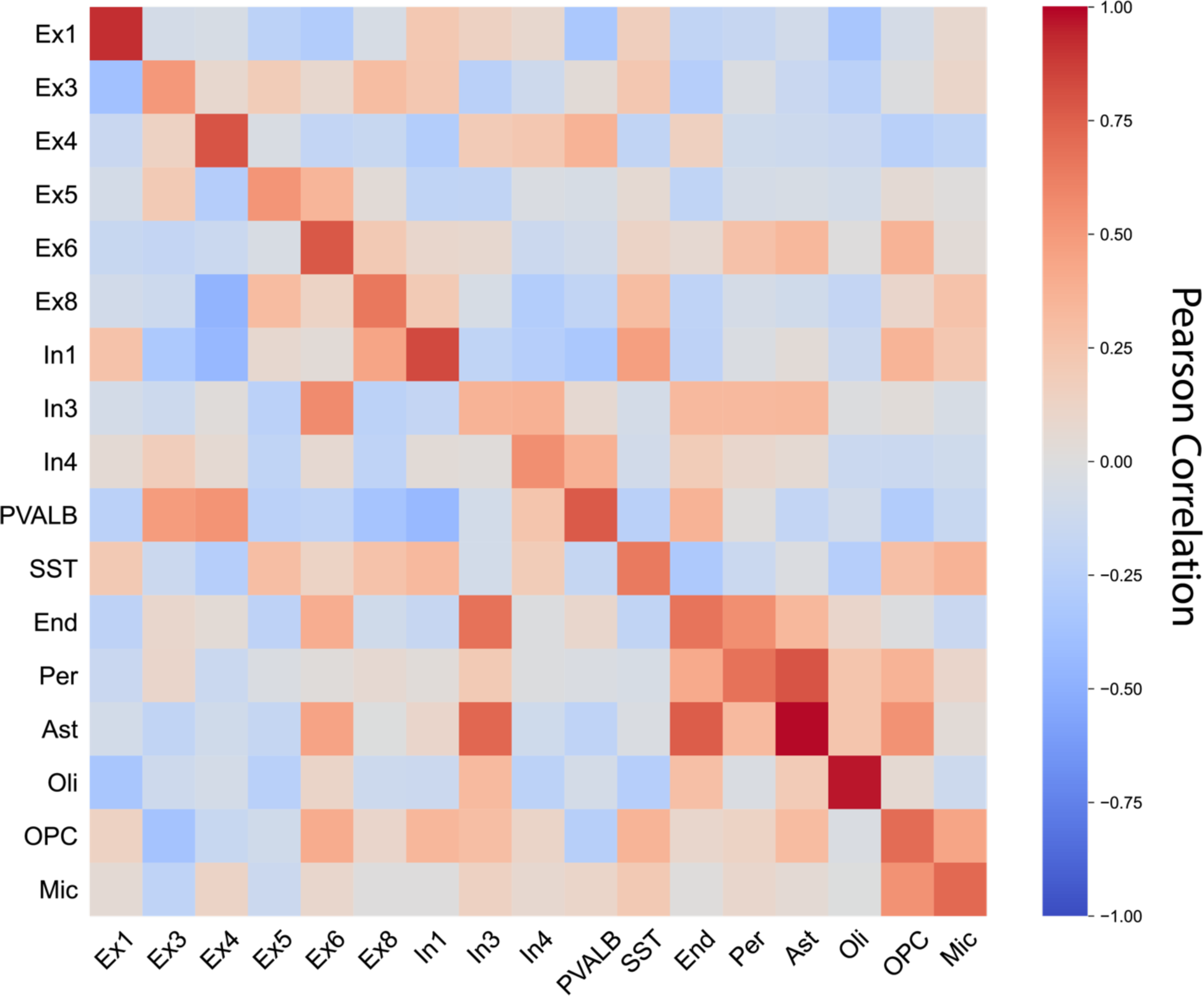
Spatial correlation of cortical cell type abundances. The abundance of each cell type across cortical parcels is correlated with other cell types. A higher correlation (in red) indicates more similar spatial distribution between cell types. Elements in the diagonal represent the correlation between the same cell type imputed from Lake DFC and Lake VIS. Elements in the upper triangle represent the correlation between two different cell types imputed from Lake DFC and the lower triangle are the correlations imputed from Lake VIS.

**Supplementary Fig 5.**
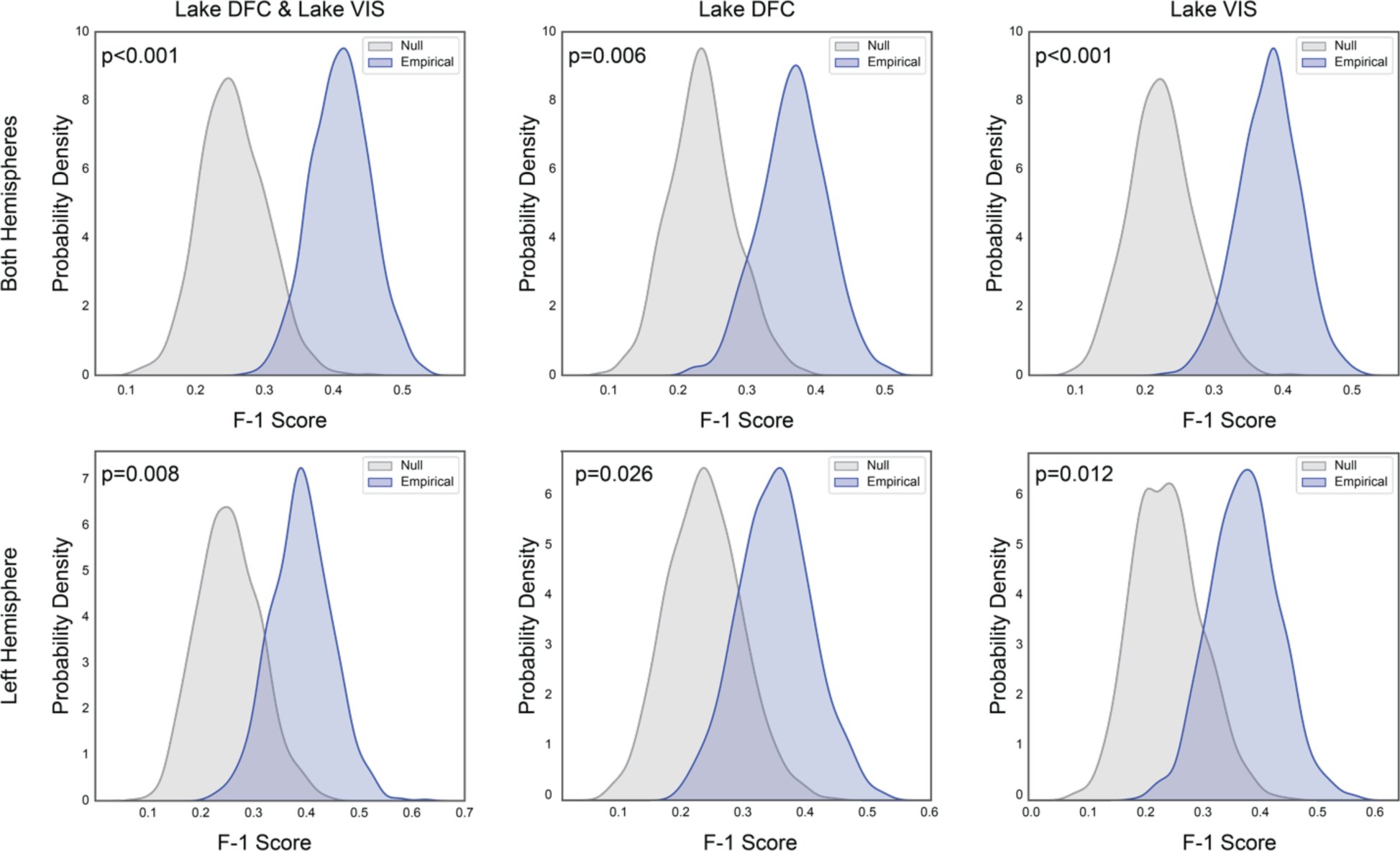
F-1 scores for models trained and tested in the six different groups. Distributions in blue were constructed from 1000 classifiers trained on real network labels, and the gray colored distribution represent classifiers trained on network labels shuffled by a Hungarian spinning method that controls for spatial autocorrelation. P-values were calculated as the percentage of null F-1 scores that are greater than the median of the empirical F-1 scores. The first row are the models trained and tested from parcels of both hemispheres, and the second row are the models trained and tested from parcels of the left hemisphere only. The first column is the models assembling information from both Lake DFC and Lake VIS. The second and the third column are the models trained from Lake DFC and Lake VIS only. Each one of the 6 ways of training and testing models yield a significant p-value for F-1 score.

**Supplementary Fig 6.**
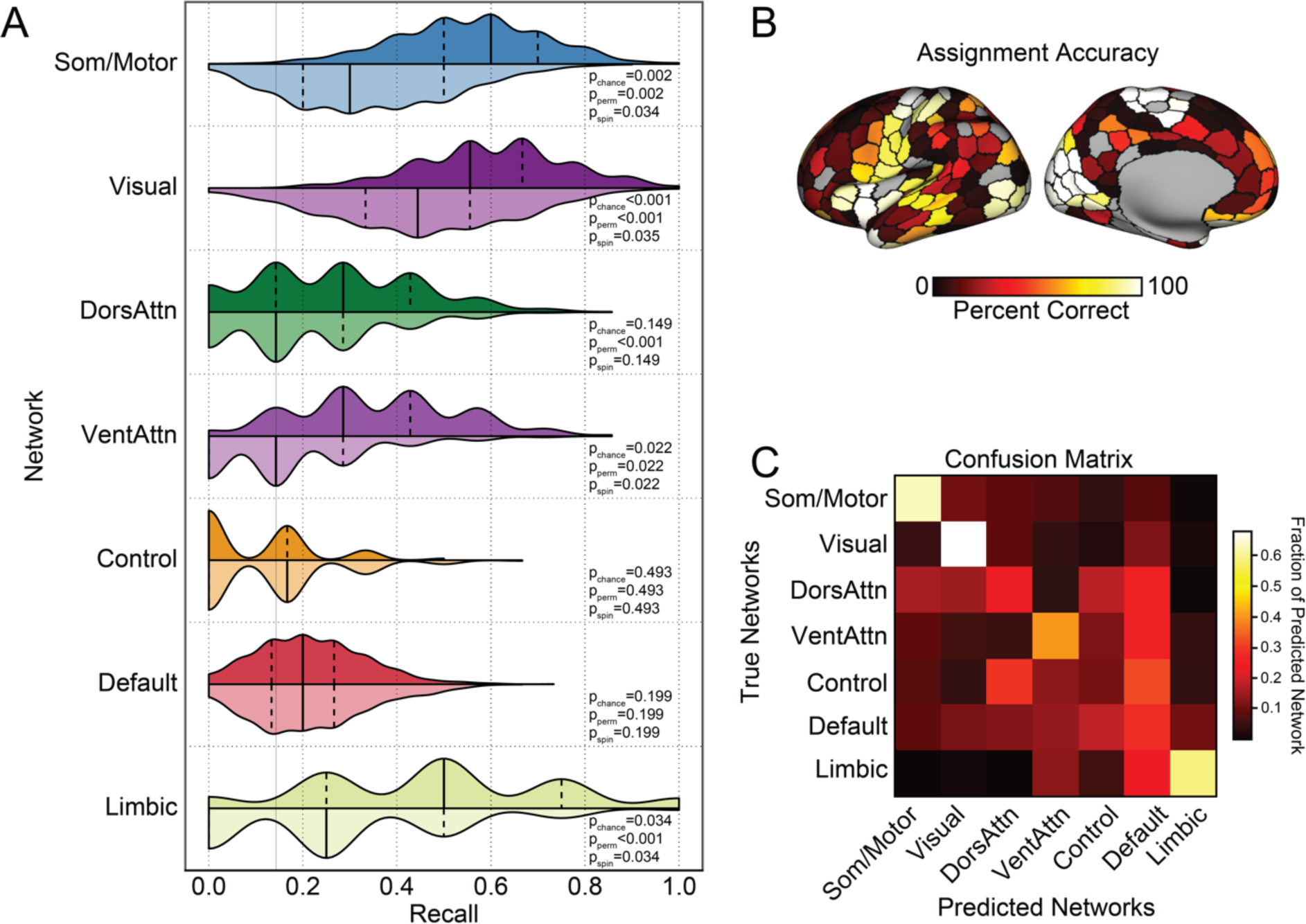
Trained and predicted on the left hemisphere only, ensembling Lake DFC and Lake VIS: Large-scale functional network assignment can be predicted by cell-type fractions in post-mortem tissue. **A.** Histograms display the SVM recall, or the probability of correctly classifying a parcel to the associated network. Consisting with Fig. 5, these data suggest the classifiers were able to predict somato/motor, visual, ventral attention, and limbic networks significantly above chance. Distributions in darker color were constructed from 1000 classifiers trained on real network labels, and the lighter colored distribution represent classifiers trained on network labels shuffled by spin-test that controls for spatial autocorrelation. The solid lines indicate median and the dashed lines represent quartiles of the distribution. **B.** Accuracy for network assignment across cortical parcels, calculated from all testing sets. **C.** Each row of the confusion matrix represents the fraction of parcels within the specific network that were predicted as belonging to each of the 7 networks. The diagonal represents the percentage of correctly classified parcels within each network. Here, the confusion matrix suggests a preferentially distinct cellular profile for somato/motor, visual, and limbic networks. While classification accuracies were low for the remaining association cortex networks, dorsal attention, default, and control networks display a higher rate of misclassifications among each other, relative to unimodal networks.

**Supplementary Fig 7.**
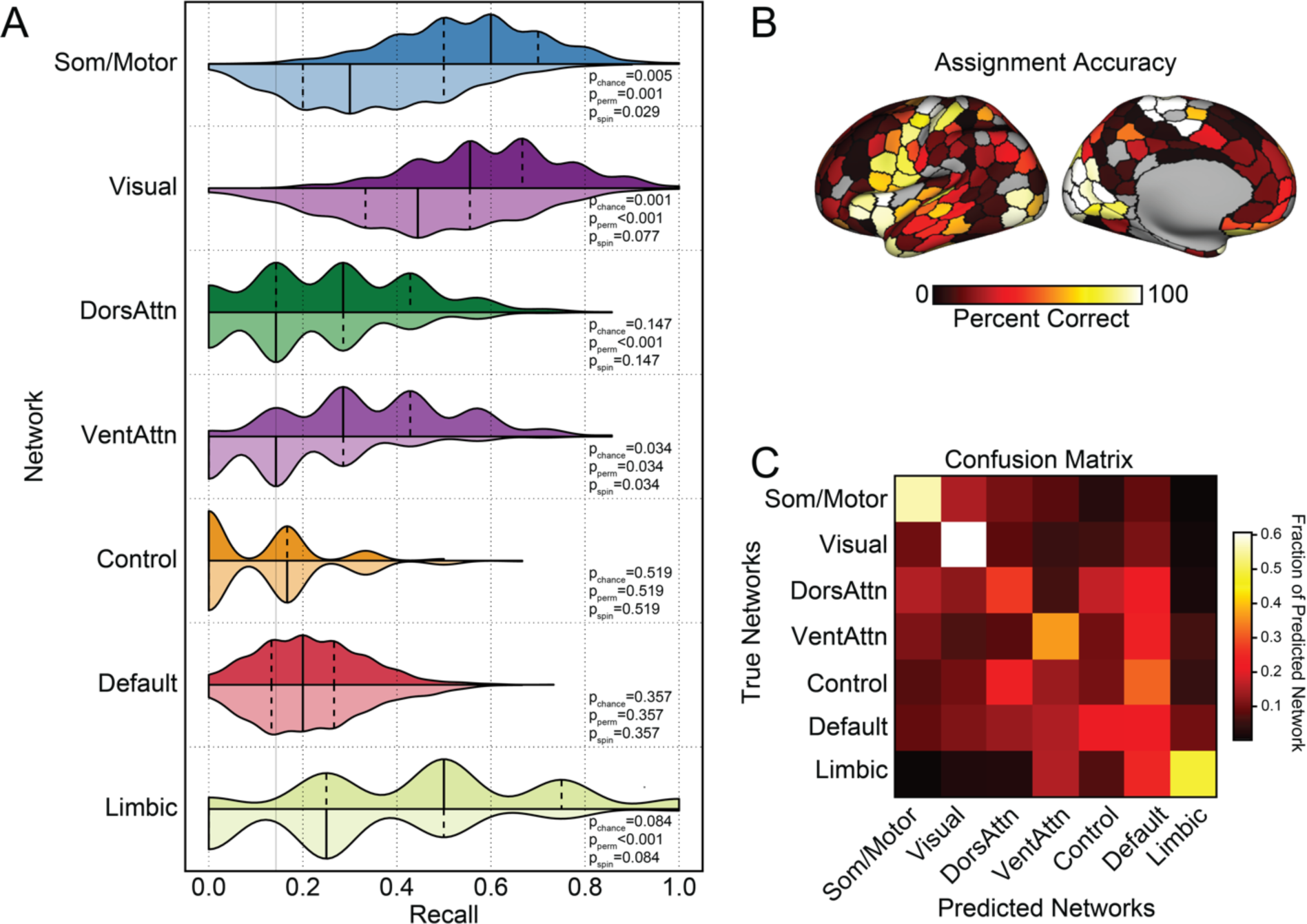
Trained and predicted on the left hemisphere only, using Lake DFC: Large-scale functional network assignment can be predicted by cell-type fractions in post-mortem tissue. **A.** Histograms display the SVM recall, or the probability of correctly classifying a parcel to the associated network. These data suggest the classifiers were able to predict somato/motor and ventral attention networks significantly above chance. Distributions in darker color were constructed from 1000 classifiers trained on real network labels, and the lighter colored distribution represent classifiers trained on network labels shuffled by a Hungarian spinning method that controls for spatial autocorrelation. The solid lines indicate median and the dashed lines represent quartiles of the distribution. **B.** Accuracy for network assignment across cortical parcels, calculated from all testing sets. **C.** Each row of the confusion matrix represents the fraction of parcels within the specific network that were predicted as belonging to each of the 7 networks. The diagonal represents the fraction of correctly classified parcels within each network. Consisting with Fig. 5, the confusion matrix suggests a preferentially distinct cellular profile for somato/motor, visual, and limbic networks. While classification accuracies were low for the remaining association cortex networks, dorsal attention, default, and control networks display a higher rate of misclassifications among each other, relative to unimodal networks.

**Supplementary Fig 8.**
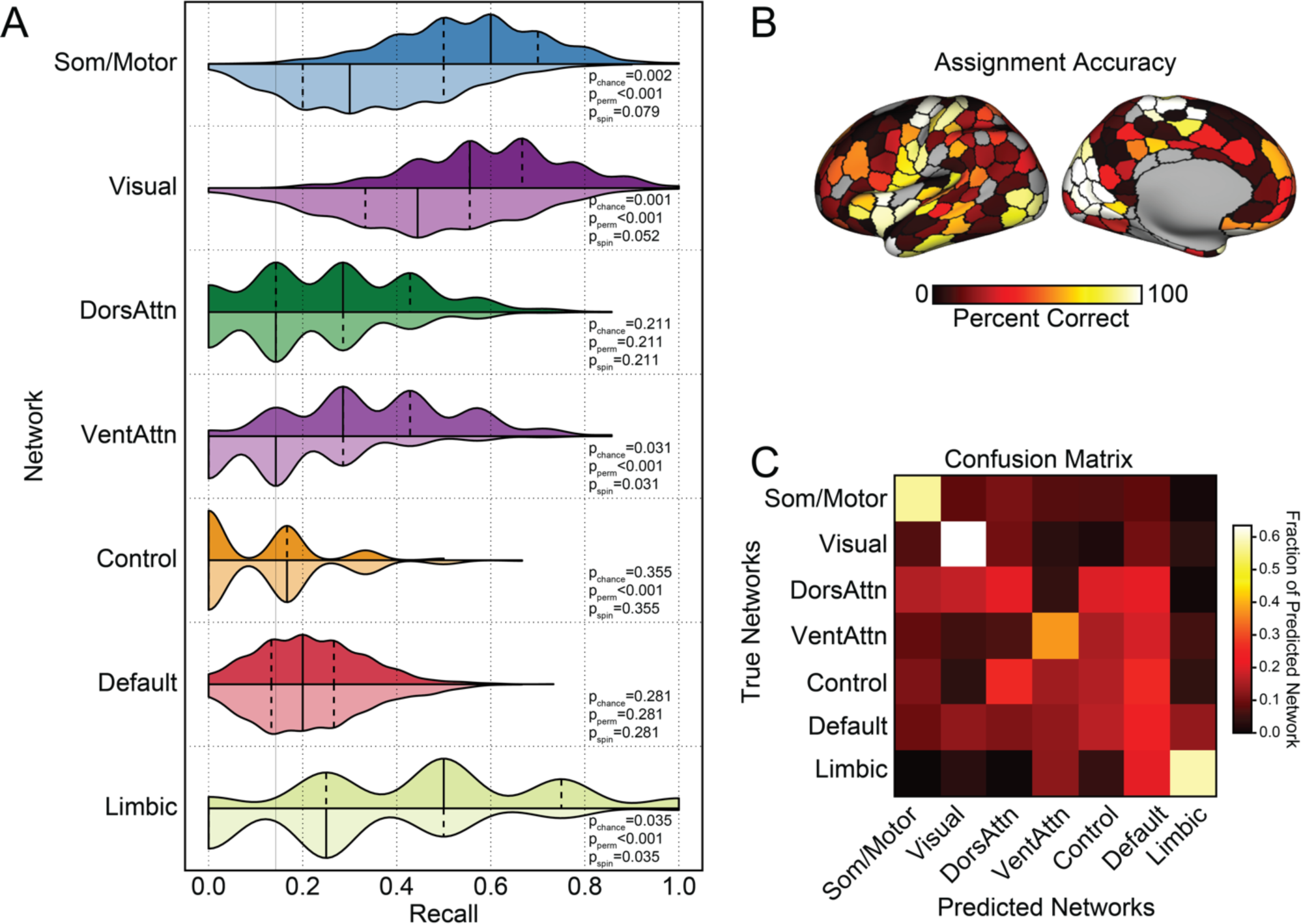
Trained and predicted on the left hemisphere only, using Lake VIS: Large-scale functional network assignment can be predicted by cell-type fractions in post-mortem tissue. **A.** Histograms display the SVM recall, or the probability of correctly classifying a parcel to the associated network. These data suggest the classifiers were able to predict ventral attention and limbic networks significantly above chance. Distributions in darker color were constructed from 1000 classifiers trained on real network labels, and the lighter colored distribution represent classifiers trained on network labels shuffled by a Hungarian spinning method that controls for spatial autocorrelation. The solid lines indicate median and the dashed lines represent quartiles of the distribution. **B.** Accuracy for network assignment across cortical parcels, calculated from all testing sets. **C.** Each row of the confusion matrix represents the fraction of parcels within the specific network that were predicted as belonging to each of the 7 networks. The diagonal represents the fraction of correctly classified parcels within each network. Consisting with Fig. 5, the confusion matrix suggests a preferentially distinct cellular profile for somato/motor, visual, and limbic networks. While classification accuracies were low for the remaining association cortex networks, dorsal attention, default, and control networks display a higher rate of misclassifications among each other, relative to unimodal networks.

**Supplementary Fig 9.**
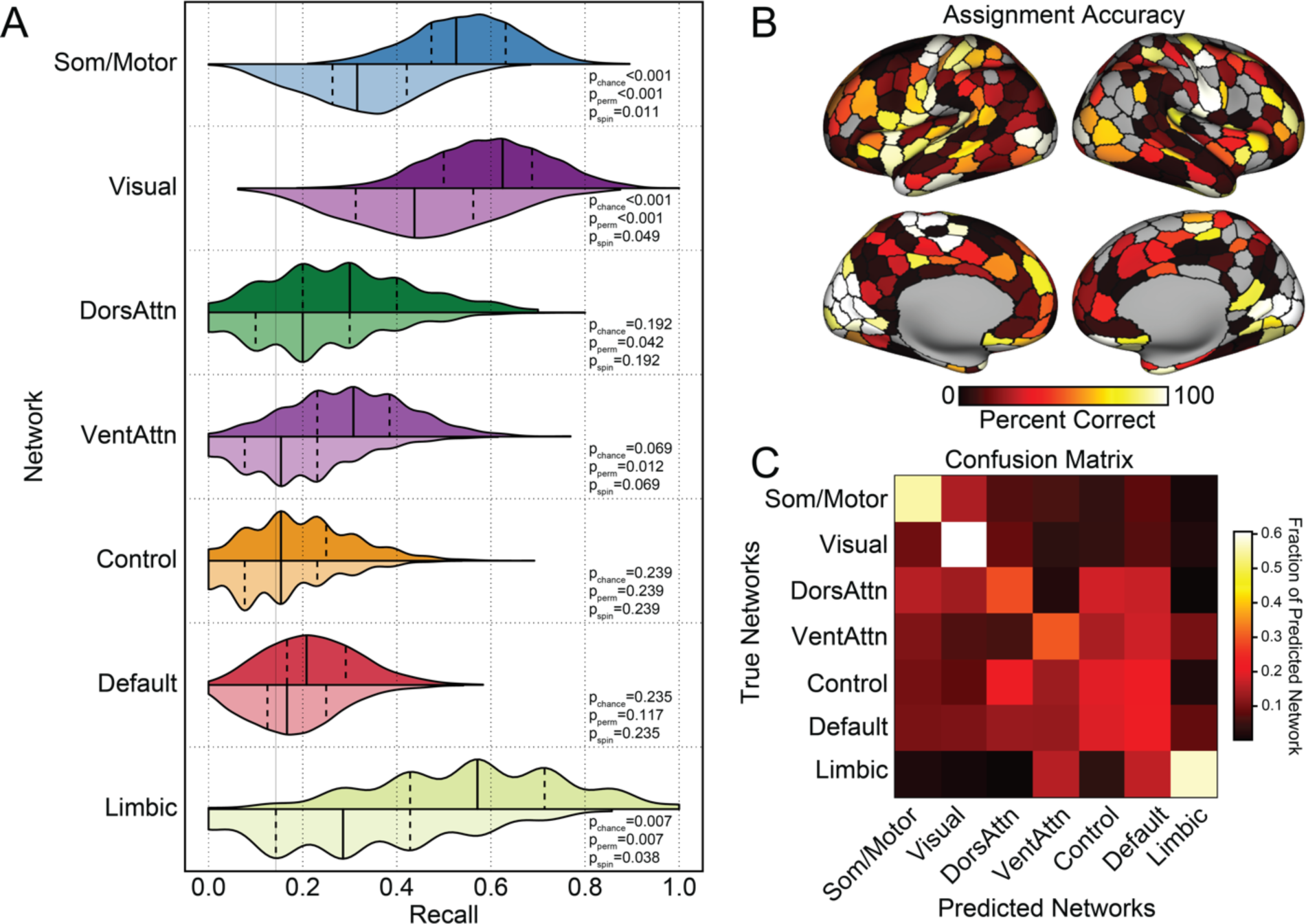
Trained and predicted on both hemispheres, using Lake DFC: Large-scale functional network assignment can be predicted by cell-type fractions in post-mortem tissue. **A.** Histograms display the SVM recall, or the probability of correctly classifying a parcel to the associated network. These data suggest the classifiers were able to predict somato/motor, visual, and limbic networks significantly above chance. Distributions in darker color were constructed from 1000 classifiers trained on real network labels, and the lighter colored distribution represent classifiers trained on network labels shuffled by a Hungarian spinning method that controls for spatial autocorrelation. The solid lines indicate median and the dashed lines represent quartiles of the distribution. **B.** Accuracy for network assignment across cortical parcels, calculated from all testing sets. **C.** Each row of the confusion matrix represents the fraction of parcels within the specific network that were predicted as belonging to each of the 7 networks. The diagonal represents the percentage of correctly classified parcels within each network. Consisting with Fig. 5, the confusion matrix suggests a preferentially distinct cellular profile for somato/motor, visual, and limbic networks. While classification accuracies were low for the remaining association cortex networks, dorsal attention, default, and control networks display a higher rate of misclassifications among each other, relative to unimodal networks.

**Supplementary Fig 10.**
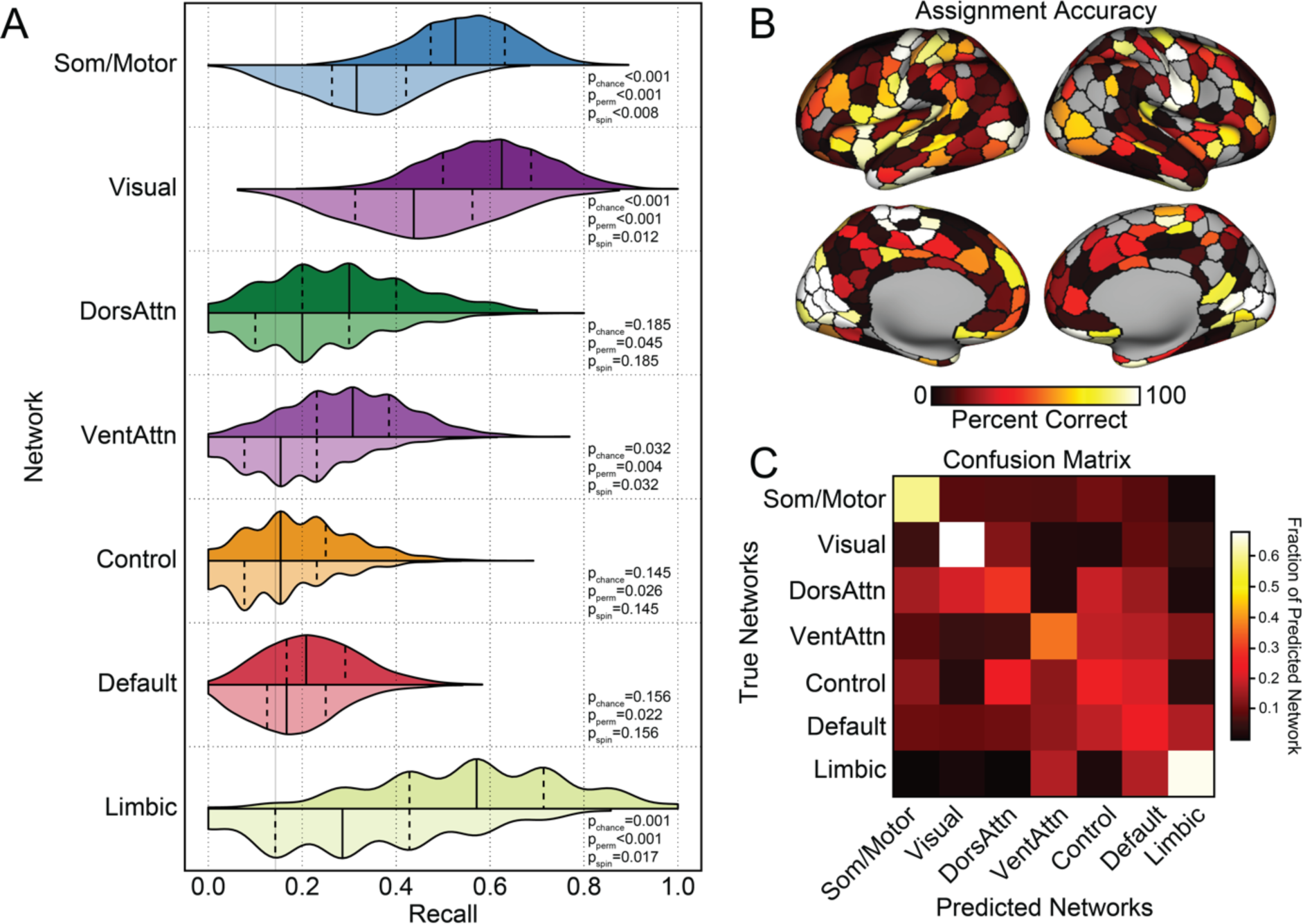
Trained and predicted on both hemispheres, using Lake VIS: Large-scale functional network assignment can be predicted by cell-type fractions in post-mortem tissue. **A.** Histograms display the SVM recall, or the probability of correctly classifying a parcel to the associated network. Consisting with Fig. 5, these data suggest the classifiers were able to predict somato/motor, visual, ventral attention, and limbic networks significantly above chance. Distributions in darker color were constructed from 1000 classifiers trained on real network labels, and the lighter colored distribution represent classifiers trained on network labels shuffled by a Hungarian spinning method that controls for spatial autocorrelation. The solid lines indicate median and the dashed lines represent quartiles of the distribution. **B.** Accuracy for network assignment across cortical parcels, calculated from all testing sets. **C.** Each row of the confusion matrix represents the fraction of parcels within the specific network that were predicted as belonging to each of the 7 networks. The diagonal represents the percentage of correctly classified parcels within each network. The confusion matrix suggests a preferentially distinct cellular profile for somato/motor, visual, and limbic networks. While classification accuracies were low for the remaining association cortex networks, dorsal attention, default, and control networks display a higher rate of misclassifications among each other, relative to unimodal networks.

